# To integrate or not to integrate: Testing degenerate strategies for solving an accumulation of perceptual evidence decision-making task

**DOI:** 10.1101/2024.08.21.609064

**Authors:** Charles D. Kopec, Thomas Zhihao Luo, Adrian G. Bondy, Diksha Gupta, Verity A. Elliott, Julie A. Charlton, Jessica R. Breda, Wynne M. Stagnaro, Emily J. Reyes, Adrian I. Sirko, Andres F. Bustos, Jovanna M. Willock, Jessica M. Morrison, Klaus L. Osorio, Carlos D. Brody

## Abstract

A common approach in the study of cognition is to train subjects to perform a task that requires a particular cognitive process to solve. Analysis of the subjects’ response behavior while they perform these tasks can offer valuable insight into the underlying mechanisms that give rise to cognition. However, if subjects are able to accurately perform such a task by using a strategy that doesn’t involve the targeted cognitive process, data from those experiments becomes more difficult to interpret. A number of perceptual decision-making tasks have been designed to study the accumulation of evidence, i.e. how noisy information presented over time is used to form a decision. Recent work, however, has highlighted how a variety of non-integration strategies can by some measures yield strikingly near-optimal performance on such tasks, raising the possibility that past conclusions from these experiments may be incorrect. Here we assemble the largest data set of animals performing one such task – the “Poisson Clicks” task – which is optimally solved by the gradual integration of pulsatile auditory noise. To investigate whether rats are in fact using this strategy, we compiled data from 515 rats performing over 35 million trials. We compare performance of 3 degenerate strategies (that circumvent the need to integrate evidence) to the optimal (integration) strategy. We demonstrate that the pulsatile nature of the stimuli used in the Poisson Clicks Task makes it possible to distinguish which strategy subjects use. Overwhelmingly, we find the rats are using an integration strategy when performing the Poisson Clicks Task.

## Introduction

“All models are wrong, but some are useful,” George Box famously opined in 1976 (Box, 1976) yet many of us often need reminding of it. The purpose of science is to improve our understanding of the natural world. We do this by constructing simplified models of the complex systems we study. The goal of these models is to make testable predictions which in turn can lead to better models and a deeper understanding of the system being studied. It’s not enough that a model can accurately describe a system, if the premise on which the model is built is fundamentally wrong. For example, the geocentric model of the solar system with its nested epicycles was able to accurately predict the location of celestial bodies in the sky (Ptolemy, 150), yet today we know this is not how the solar system functions (Copernicus, 1543). Therefore we must constantly strive to test our models and be willing to toss them out when necessary (Palminteri et al., 2017; Wilson and Collins, 2019).

In neuroscience we seek to understand how the brain functions, from how the intricate interactions of molecules give rise to cells, to how the coordinated activity of those cells give rise to behavior and cognition. A common strategy in studying cognition is to train subjects to perform a task that requires a particular cognitive process to solve accurately (Thorndike, 1911). We can then build mathematical models to describe the subjects’ behavior with the goal of better understanding the underlying cognitive phenomenon (Bogacz et al., 2006; Wilson and Collins, 2019). Here again George Box provides insight: “Since all models are wrong the scientist must be alert to what is importantly wrong. It is inappropriate to be concerned about safety from mice when there are tigers abroad.”(Box, 1976). If it is possible to accurately perform a task without employing the cognitive process it’s designed to test (Ditterich, 2006; Stine et al., 2020), the models that assume its engagement should be questioned. For example, if a subject could write down the list of words in a short-term memory task and then later when asked to recite them simply read back what they had written, modeling performance of this task would teach us little about memory. We must be vigilant to the possibility that subjects are using alternative strategies to perform the tasks we train them on, and run additional tests and analysis to show they are not (Glickman and Usher, 2019; Hyafil et al., 2023; Pinto et al., 2018).

The world we inhabit is complex, and sensory information that reaches us is often corrupted by noise. To get a better sense of what one is hearing, seeing, smelling, or feeling, one can reduce this noise by averaging over multiple independent samples. The accumulation of sensory evidence is an integral component of perceptual decision making. How the brain is able to combine noisy information presented gradually over time and use that to form a decision of what action to take has been the focus of extensive study for many decades (Brody and Hanks, 2016; Hanks and Summerfield, 2017; Newsome and Paré, 1988). To understand this process better, cognitive neuroscientists have created a collection of perceptual decision-making tasks that target different sensory modalities including audition (Brunton et al., 2013; Hanks et al., 2015; Pardo-Vazquez et al., 2019; Raposo et al., 2012; Sanders and Kepecs, 2012), vision (Gold and Shadlen, 2000; Katz et al., 2016; Morcos and Harvey, 2016; Odoemene et al., 2018; Raposo et al., 2012; Scott et al., 2015; Shadlen and Newsome, 1996), olfaction (Bowman et al., 2012; Uchida et al., 2006), and somatosensation (Deverett et al., 2018), and applied them to a range of species from rats (Brunton et al., 2013; Hanks et al., 2015; Pardo-Vazquez et al., 2019; Raposo et al., 2012; Scott et al., 2015) and mice (Deverett et al., 2018; Morcos and Harvey, 2016; Odoemene et al., 2018; Pinto et al., 2018; Sanders and Kepecs, 2012), to monkeys (Gold and Shadlen, 2000; Katz et al., 2016; Shadlen and Newsome, 1996) and humans (Brunton et al., 2013; Cheadle et al., 2014; Glickman and Usher, 2019; Keung et al., 2019; Raposo et al., 2012; Wyart et al., 2012). In these tasks, subjects are typically presented with a stream of noisy sensory stimuli, they must attend to some component of that stimulus, and use it to form a binary decision. For example, in the Random Dot Motion Task (Gold and Shadlen, 2000; Newsome and Paré, 1988) subjects are shown a field of moving dots, with some moving in a coherent direction and the rest moving randomly. The subject is rewarded if they correctly report the direction of coherent motion. In the Poisson Clicks Task (Brunton et al., 2013; Sanders and Kepecs, 2012) subjects are played two streams of randomly timed auditory clicks, one coming from a speaker to their right and the other from a speaker to their left. Here, the subject is rewarded if they correctly report which speaker played more clicks.

One additional feature of perceptual decision-making tasks is how the stimuli are presented to the subject. Tasks can generally be categorized into two groups. The first are continuous evidence tasks, this includes the Random Dot Motion Task (Gold and Shadlen, 2000; Newsome and Paré, 1988; Shadlen and Newsome, 1996) as well as the Cloud of Tones Task (Znamenskiy and Zador, 2013), where sensory evidence is presented continuously to the subject. Continuous evidence tasks have no pauses or gaps in the sensory stream and multiple bits of evidence are presented simultaneously. The other group are discrete evidence tasks. These include the Poisson Clicks Task (Brunton et al., 2013; Sanders and Kepecs, 2012) as well as a visual version known as Flicker (Scott et al., 2015), a spatial navigation version dubbed Accumulating Towers (Harvey et al., 2012; Pinto et al., 2018), and many others (Bronfman et al., 2015; Cheadle et al., 2014; Cisek et al., 2009; Morillon et al., 2014; Pardo-Vazquez et al., 2019; Wyart et al., 2012; Yang and Shadlen, 2007) where the sensory stimulus is composed of discrete bits of information often separated by periods of no information. Hybrid tasks also exist between these two categories where a series of short discrete periods of continuous evidence are presented to a subject (Levi et al., 2023; Yates et al., 2017). This particular, seemingly nuanced feature of perceptual decision-making tasks will be highly relevant to the analysis we present here. However, regardless of how the stimuli are presented, the optimal strategy in all of these tasks is the same, and requires the subject to integrate over the entire stimulus. This is because at any individual moment in time the stimulus is too noisy to be accurately relied upon.

The models used to describe the decision-making behavior in these tasks are often built around a drift-diffusion mechanism (Bogacz et al., 2006; Bronfman et al., 2015; Brunton et al., 2013; Busemeyer and Townsend, 1993; Cisek et al., 2009; Gold and Shadlen, 2007; Ratcliff, 1978; Ratcliff and McKoon, 2008) and generally assume the subject is integrating the entire stimulus. However, if subjects are not (Thura et al., 2012), then this is a tiger George Box warned about, making the tasks, models, and the conclusions we draw from them less reliable (Hyafil et al., 2023; Stine et al., 2020). Here lies the important question: What strategy are subjects using to perform accumulation of perceptual evidence decision-making tasks? While there are strategies subjects can use to perform reasonably well on these tasks without integrating the entire stimulus, simple analysis of their behavioral responses is usually sufficient to demonstrate subjects do not tend to use them. One approach to determine a subject’s strategy is to develop more universal models that are able to adopt a range of different strategies (Wilson and Collins, 2019), or a series of different models that each assume a unique strategy; and determine which better explains the subjects’ response behavior. Previous work on the Poisson Clicks Task used a model that was able to account for a few different suboptimal strategies (Brunton et al., 2013), including a burst detection strategy, which we will explore further here. The overall conclusion from this previous work was that rats integrate the entire stimulus in a nearly optimal way. However, this model still left many strategies untested.

Recent work (Stine et al., 2020) investigating monkey and human performance on the continuous evidence Random Dot Motion Task found that simple behavioral analyses and model fitting were not sufficient to determine a subject’s strategy. Specifically, they identified two strategies, dubbed Extrema (referred to here as Burst) and Snapshot (see also (Cartwright and Festinger, 1943; Ditterich, 2006)), where a subject does not integrate the entire stimulus, but is still able to generate responses that make it appear as if they are. Here we refer to this class of strategies, where the subject does not use the cognitive process the task is designed to probe, yet is still able to generate responses appearing as if they do, as degenerate strategies. The conclusion of this study was that only modifications to the accumulation of evidence task structure could allow one to determine if a subject was integrating the entire stimulus or not. They found that only when subjects perform both trials with a variable stimulus duration and trials where the subject is free to respond at any time during the stimulus could they conclude if the entire stimulus was being integrated.

More recent work however has challenged this conclusion (Hyafil et al., 2023). Investigating monkey, human, and rodent performance on a series of different discrete evidence accumulation tasks, the authors conclude that analyses do in fact exist which allow one to determine if a subject is using one of the degenerate strategies identified by Stine et al., or integrating the entire stimulus, without any alteration of the task structure. Here we will use one of the analyses they identified, which leverages “disagree trials” (Hyafil et al., 2023; Levi et al., 2018) (we refer to these as “conflict trials”), on behavioral data from rats performing the Poisson Clicks Task. The strategy a subject uses to perform a task can leave a particularly strong fingerprint on the pattern of mistakes they make. Different strategies predict different patterns. Identifying a strategy that predicts a similar pattern of mistakes to what the subject produces is strong evidence the subject is using that strategy. Here we make use of conflict trials as one test to identify which strategy rats use to perform the Poisson Clicks Task. We further identify additional novel analysis specifically tailored to discrete evidence tasks like the Poisson Clicks Task, and when applied, consistently find evidence that rats integrate information from the entire stimulus.

In this study, we will first test if degenerate strategies can be used to successfully perform a discrete evidence auditory perceptual decision-making task, the Poisson Clicks Task, in a way that mimics a subject integrating evidence from the entire stimulus, i.e. full integration. Second, we will demonstrate that it is possible, given the task structure, to determine whether or not these degenerate strategies are being used. The dataset on which we will test these strategies comes from 515 individual male rats performing a collective 35,858,295 trials (Kopec et al., 2024). We expand the original model used to describe behavior on the Poisson Clicks Task (Brunton et al., 2013) and develop a series of degenerate strategy models, to more fully explore how rats perform this task. We will first test the two strategies identified by Stine et al. using a combination of model-free analysis and model fitting: 1) Burst (Ditterich, 2006; Stine et al., 2020; Waskom and Kiani, 2018), where the subject only responds to a brief burst of strong evidence; 2) Snapshot (Pinto et al., 2018; Stine et al., 2020), where the subject integrates only a very small portion of the entire stimulus. Subsequently, we will test a third degenerate strategy that combines elements from these first two: 3) Single Click, where the subject responds to only a single randomly selected click. Overall, we demonstrate that it is possible to use models (Wilson and Collins, 2019) and model free analysis (Palminteri et al., 2017) to determine whether or not subjects are using one of these degenerate strategies, without altering the task structure. We conclude that the vast majority of subjects in this dataset are not (99.8% of rats are likely not using a burst strategy, 88.9% are likely not using a snapshot strategy (those that may be are using a very long snapshot nearly equivalent to full integration), and 100% are likely not using a single click strategy).

The analyses performed here, and the ability to make the determination that rats are not using these degenerate strategies, rely heavily on the discrete pulsatile nature and Poisson statistics of the stimulus used in the Poisson Clicks Task. The potential to analyze existing data and determine a subject’s strategy without having to alter the task structure and collect new data, highlights a significant advantage discrete evidence tasks have over continuous evidence accumulation of evidence tasks. The findings presented here demonstrate that the body of literature, relying on the assumption subjects in the Poisson Clicks and similar tasks integrate information from the entire stimulus, does not suffer from a crucial flaw suggested by Stine et al. Finally, we conclude there is likely no conflict between the findings from Stine et al. which focused on continuous evidence tasks and Hyafil et al. which focused on discrete evidence tasks; rather their conclusions were driven by the limitations of the analyses imposed by the different stimuli used in the tasks they investigated.

## Results

### The Poisson Clicks Task and Dataset

We have compiled data from 515 fully trained rats performing over 35 million behavioral trials on the Poisson Clicks Task. The data was collected from 135,326 sessions (continuous periods of time in an operant behavior chamber), with each session averaging 128 minutes, performed over a 15 year period. Collectively this data represents 33 years of behavioral data. In this work, we use this dataset to test various strategies rats may be using to perform the Poisson Clicks Task. To facilitate study by others we’ve made this dataset available on zenodo.org (Kopec et al., 2024)(see Methods: Dataset).

The Poisson Clicks Task is an auditory perceptual decision-making task designed to study the gradual accumulation of information (Brunton et al., 2013). The task structure is outlined in Figure 1a, and performed in an operant chamber where rats have access to 3 nose ports, cylindrical recesses where they can poke their nose (see Methods). Trials are self initiated by the rat who is cued with a LED in the center port. When they are ready to perform a trial the rat pokes his nose into the center port and holds it there. After a variable delay of silence two streams of auditory clicks play from each of two speakers positioned above the left and right nose ports. In the “location” variant of the task, each speaker plays an independent stream of randomly timed white noise clicks. In this version of the task the first click is often a stereo click played simultaneously from both speakers. When the clicks stop playing the center LED is extinguished and the rat is free to withdraw from the center port. The rat is rewarded with a small drop of water if they poke their nose into the side port associated with the speaker that played the greater number of clicks. An incorrect response is paired with a burst of white noise and a timeout before the next trial can begin. If the rat fails to hold its nose in the center port during the entire fixation period the trial is aborted as a violation and a new trial begins after a short delay. Violation trials are excluded from all analysis in this study.

**Figure 1.**
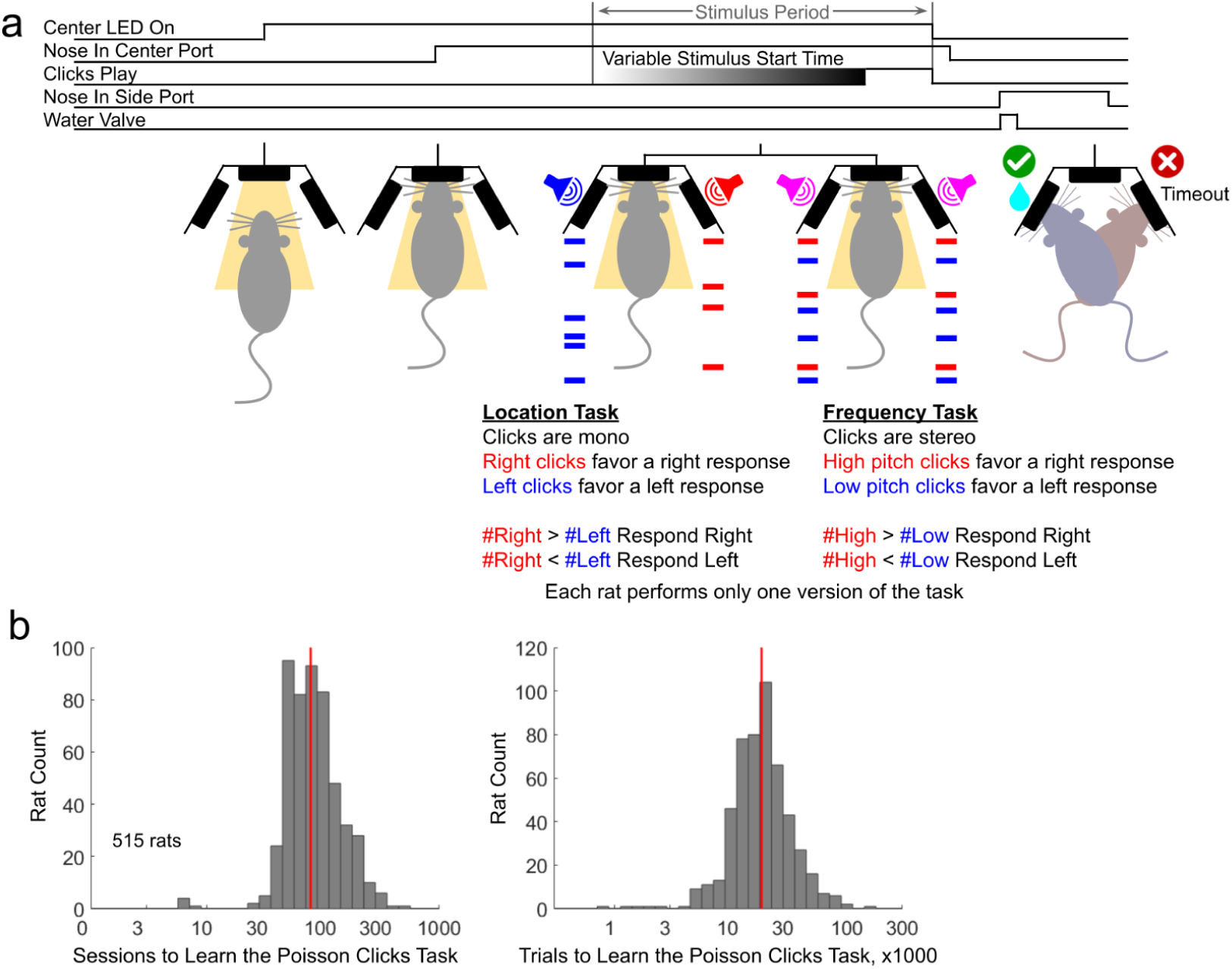
The Poisson Clicks Task. **a)** Poisson Clicks task structure. The center port light turning on is the cue to the subjects that they are free to begin a trial. When ready, they poke their nose into the center port and are trained to hold it there under the center light is turned off. While the animal has its nose in the center port, after a variable duration silence, two streams of randomly timed clicks are played from speakers above each side port. The stimulus plays for a variable duration. When the clicks finish playing, the center light is extinguished and the subject is free to withdraw from the center port. A subsequent poke into the correct side port provides a small water reward, while an incorrect side poke is paired with a white noise sound and a time out before the next trial can begin. Rats are trained on one of two different versions of the Poisson Clicks Task. In the Location version of the task, each click is played from either the left or right speaker, and the correct side is the one that played the greater total number of clicks. In the Location version the first click is often a stereo click played from both speakers simultaneously as depicted here. In the Frequency version of the task, all clicks are played in stereo, and each click is either a low or a high tone. The correct side is Right if more high tones were played, and Left if more low tones were played, **b)** Histograms of the number of training sessions (left) and trials (right) the rats required before mastering the Poisson Clicks task, with mastering defined as the time from first introduction to a training rig, until reaching performance criteria sufficient to being included in this study (see main text for criteria). Medians are marked with a red line (78 sessions, 18,379 trials). Note the log scale of the horizontal axis.

An alternate “frequency” version of the task all clicks are stereo and consist of a 3 ms pure tone, either low 6.5 kHz or high 14.2 kHz. Here the rat must decide if more high or low tone clicks were played. In the frequency version high and low clicks are never allowed to overlap, and therefore there is no equivalent of an initial stereo click as in the location version. Rats are only trained on one version of the task. Throughout this study data from both versions of the task are combined as we find no significant difference in performance between them.

The difficulty of individual trials can be controlled by adjusting the Poisson rate of the “left evidence” and “right evidence” streams. Typically reported as gamma, this is the log ratio of the right and left evidence rates:

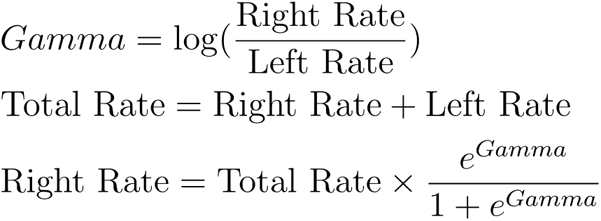

The stimulus period is defined as the time when clicks may play. This is typically 1 s long beginning 500 ms after the rat pokes into the center port and terminates when the center port LED is extinguished (Figure 1a). The total duration of the stimulus is variable and ranges between 200 ms and 1 s.

Typical trials on the Poisson Clicks Task are generated using a unique seed in a pseudorandom list, generating an entirely unique stimulus. The random timing of the clicks allows us to better sample the stimulus space and constrain model fits. However, while a model will yield a probability of producing a particular response, the subject either does or does not produce that response only once following that stimulus. One can maximize over the probability of the model matching the subject’s responses to constrain model parameters, but we still wondered, if a model predicts that a subject will produce a rightward response some fraction of the time, say 70%, if we repeated that trial to the subject multiple times, say 100 replicates, would we in fact see responses in line with the model prediction, i.e. 70 right and 30 left choices. These repeated stimulus trials, dubbed “frozen noise” trials, are randomly presented to the rats and generally make up no more than 50% of the trials. Typically 10 different trials per generative gamma are presented; given 8 generative gamma values +/- [0.5, 1.5, 2.5, 3.5] for a total of 80 different frozen trials. A rat will experience each frozen noise trial on average 2.5 times given a typical session of 400 trials. Not all rats trained on the Poisson Clicks Task experienced frozen noise trials. For each of the strategies tested in this study we use the frozen noise trials as a held out test to see how well the different strategies can predict the rats’ performance. The models are fit to the unique (non-frozen) trials only and then run on the frozen noise trials with their predicted performance plotted against the rats’ response behavior (see Figures 4b, 6b, 8b, and 9e below).

The dataset in this study is pooled from all rats in the Brody lab that have trained on the Poisson Clicks Task (“location” and “frequency” versions) and meet the following criteria: 1) perform the task for a minimum of 100 days; 2) sessions (a contiguous period of time in a training rig, typically only 1 per day) must include trials of three different difficulties (absolute value of gamma); 3) the most difficult trials must have a |gamma| ≤ 1; 4) the easiest trials must have a |gamma| ≥ 3; 5) overall performance during the session must exceed 65% correct; and 6) over sessions that meet these criteria, the rat must complete a minimum of 10,000 trials (excluding violation trials). With these criteria in place, we have assembled a dataset consisting of 35,858,295 trials from 515 rats. All analysis in this study excludes the following: violation trials where the rat breaks fixation before cued, trials where an optogenetic stimulus occurs, trials with a stimulus duration greater than 1 second, “free choice” trials where no clicks are played and both choice ports are rewarded, trials where the correct response is cued with a side LED, and rats that trained for a significant period of time on a task other than Poisson Clicks. The full dataset does contain these individuals, trials, and sessions with metadata to facilitate additional study by others.

Rats learn the Poisson Clicks Task in a semi-automated training facility. Technicians load the animals into training rigs, but training occurs at the animal’s own pace as the software advances them through a series of stages shaping their behavior based on their own performance metrics (see Methods). The rats included in this data set required a median of 78 training sessions to fully learn the task (Figure 1b left, 95% of rats required between 36 and 241 sessions), defined as advancing from completely naive about the training rig to providing the first data that meets criteria defined above to be included in the dataset. This translates to a median of 18,379 trials (including violation trials) to learn the task (Figure 1b right, 95% of rats required between 5,443 and 60,932 trials)

### Rat Performance

After training rats are able to perform well on the Poisson Clicks Task as can be seen by plotting the fraction responding “right” as a function of click difference. Here we fit a 4-parameter sigmoid function to each individual rat’s data (Figure 2a, see Methods). On average, rats respond “right” only 11.7% of the time (95% of the rats are between 4.5 - 22.9%) on trials with the greatest negative click difference (i.e. when there were more left clicks than right clicks), and 87.8% of the time (95% of the rats are between 76.6 - 95.0%) on trials with the greatest positive click difference (Figure 2a panel 3, lower asymptote (upper asymptote): the easiest trials with left (right) as the correct response). Rats show minimal bias in their performance (Figure 2a panel 4), the median click count difference that results in equal fraction of left and right responses is -0.09 clicks (95% of the rats are between -4.57 clicks (biased left) to +3.42 clicks (biased right)). Finally, rats are sensitive to small changes in the overall click count as measured by the steep slope at the inflection point in the psychometric plot (Figure 2a panel 5). On average (median) rats respond “right” 3.1% more in response to 1 additional right click (95% of the rats are between 1.6 - 7.3%).

**Figure 2.**
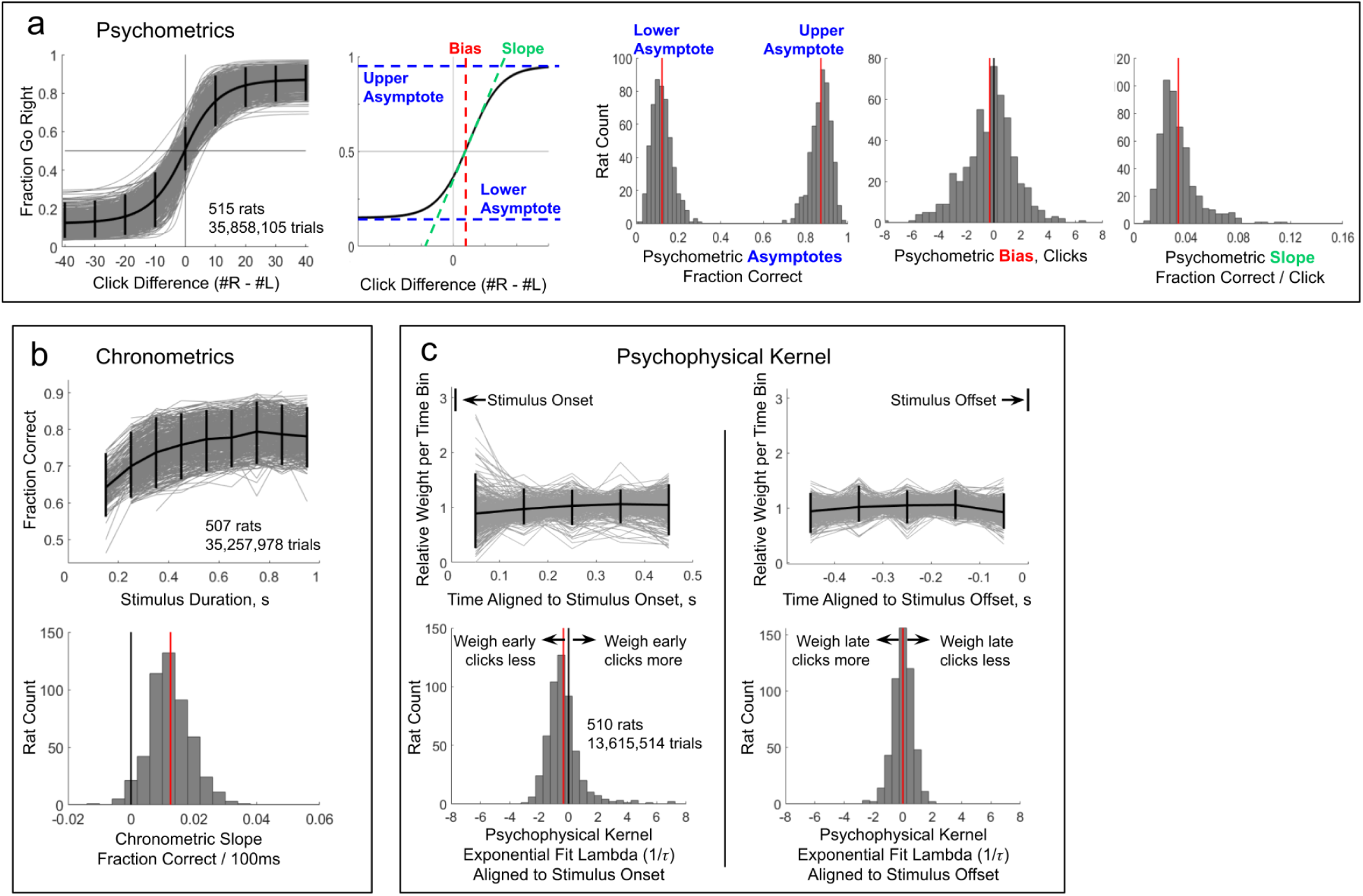
Performance of rats on the Poisson Clicks task. **a)** Left: Individual psychometric curves (fraction respond “right’’ as a function of total right minus total left clicks) plotted in gray for 515 rats. Mean psychometric curve in black. Vertical bars indicate 95% of the rats. Right: Schematic of an example psychometric curve, indicating the asymptotes, bias, and slope, followed by histograms over the individual psychometric curves of their asymptotic endpoints (lower and upper); bias (click difference that results in a Fraction Go Right = 0.5); and slope (computed at the curve’s inflection point), **b)** Upper: Individual chronometric plots (fraction correct as a function of stimulus duration) plotted in gray for 507 rats. Rats with at least 50 trials in each of at least 5 different duration bins are included in this analysis. Mean chronometric plot in black. Vertical bars indicate 95% of the rats. Lower: Histogram of the slope (fraction correct per 100 ms) of the best fit line to individual rat’s trial durations and responses. Mean marked in red. 97.0% of rats have a positive slope, p << 0.01. **c)** Upper: Psychophysical Kernel, fit to the rat’s choice data using logistic regression. This is an estimate of the influence evidence in each time bin has on the subject’s decision. Data aligned to stimulus onset (left) or stimulus offset (right). Individual rats shown in gray, mean in black. Vertical bars indicate 95% of the rats. Only trials with stimuli 500 ms or greater are included in this analysis. Lower: An exponential function is fit to each rat’s psychophysical kernel. A histogram of the lambda (A = 1/r) values from each rat’s best fit exponential curve is plotted. Mean marked in red. A = 0 indicates a flat psychophysical kernel (all clicks are influence the decision equally). Data aligned to stimulus onset (left) 72.2% rats A < 0 (p < 0.01). Aligned to offset (right) 50.0% rats A < 0 (p = 0.91).

In the Poisson Clicks Task, the reward is offered at the side associated with the stream that played more clicks (Figure 1a). Therefore, the optimal strategy is to integrate clicks from the entire stimulus to inform one’s decision. Three different analyses of the rats’ response behavior are consistent with a strategy in which they integrate the entire stimulus. First, if we plot their psychometric curves (Brunton et al., 2013; Carandini and Churchland, 2013), fraction “go right” as a function of the overall click difference (Figure 2a panel 1), we see a sigmoid monotonic relationship. More right evidence clicks leads to a greater probability of the rat responding right. Second, if we plot the chronometric curves (Brunton et al., 2013; Kiani et al., 2008; Kiani and Shadlen, 2009; Shadlen and Newsome, 1996), fraction of correct responses as a function of the stimulus duration (Figure 2b top), we see a gradual but steady increase (p < 0.01 population slope is flat). Overall, the majority of rats show an increased performance for longer stimuli (Figure 2b bottom). Third, if we compute the psychophysical kernel (see Methods), an estimate of the weight of the influence different periods within the stimulus have on the decision (Keung et al., 2020; Kiani et al., 2008; Nienborg and Cumming, 2007; Shadlen and Newsome, 1996), we see a relatively flat function, consistent with all clicks being considered equally (Figure 2c top)(Brunton et al., 2013). In Figure 2c bottom, we fit an exponential function to each rat’s psychophysical kernel and plot a histogram of the lambda values (inverse of the time constant). A lambda of zero corresponds to a flat kernel. Overall, there is a small under-weighting of the clicks that occur at the beginning of the stimulus (Figure 2c left, p < 0.01, however some rats overweight these clicks, 95% of rats’ lambda when aligned to stimulus onset is between -1.8 and 2.0) but later clicks are weighed uniformly (Figure 2c right bottom, p = 0.69, 95% of rats’ lambda when aligned to stimulus offset is between -1.6 and 1.3). Taken together these three analyses are consistent with the rats using information from the entire stimulus when deciding on a response.

A note that will be relevant for the degenerate strategies discussed below, the value of the psychophysical kernel can have different causes. On the one hand, it can refer to the weight assigned to clicks occurring at a particular time, i.e. a value of 0.5 means any click occurring at that time always adds +/- 0.5 to the accumulated total rather than +/- 1. On the other hand, a value of 0.5 can indicate that a click occurring at a particular time has a 50% probability of being sampled. When the click is sampled it adds +/- 1 to the accumulated value and when it’s not sampled it adds 0. The value of the kernel can also refer to any mixture of these two processes. A decaying kernel can either indicate a strategy that samples early clicks with a greater probability than it samples later clicks, or a strategy that samples all clicks but weighs late clicks less. When discussing the kernels resulting from the degenerate strategies we know the values refer to the probability of sampling clicks at particular times (since that’s how we designed the strategies to function). However, we must remain agnostic to the true underlying cause of the kernel fit to the rats’ data, and simply use its shape to help elucidate the strategy they’re using to perform the task.

### Degenerate Strategies

While the ideal strategy, integrating the entire stimulus, may maximize the reward earned, it is possible to use alternate strategies to perform the Poisson Clicks Task significantly better than chance. We refer to these as degenerate strategies since they allow one to perform the task reasonably well, without using the cognitive process this task is designed to probe, integration. For example, given the typical range of gamma values used to generate stimuli (+/- 0.5, 1.5, 2.5, 3.5), the first non-stereo click will indicate the correct response ∼83% of the time. However, if this click was consistently used to inform a subject’s decision, it would be clearly evident in the plots shown in Figure 2b,c. The chronometric plots (Figure 2b) would not show a change in performance as a function of stimulus duration since all clicks following the first non-stereo click would be irrelevant. The psychophysical kernel (Figure 2c) would show all the weight assigned to clicks in the first time bin and zero weight after that. A similar outcome would occur if one always relied only on the last click. Therefore, we can rule out such simple degenerate strategies given basic behavioral analysis.

Recent work has identified alternate degenerate strategies (Stine et al., 2020), dubbed Extrema (referred to here as Burst) and Snapshot, for solving a continuous evidence integration task, the Random Dot Motion Task. If a subject were to use these strategies, their response data would result in a monotonic sigmoid psychometric plot, an increasing chronometric plot, and a flat psychophysical kernel. Simple behavioral analyses are therefore not able to distinguish these strategies from full integration. The authors further demonstrate that only modifications to the task structure itself, requiring subjects to perform both variable stimulus duration trials and reaction time trials, allows one to distinguish whether the subjects are using full integration or either of these degenerate strategies to solve the task. Here, we will investigate these two degenerate strategies along with a third Single Click strategy in relation to a discrete evidence integration task, the Poisson Clicks Task. We seek to answer two questions: first, is it possible to differentiate if a subject performing the Poisson Clicks Task (as outlined in Figure 1a) is using one of these degenerate strategies or full integration; and second, which strategy are the rats in this dataset most likely using to perform the Poisson Clicks Task.

### Testing the Burst Strategy

Using the burst strategy (referred to as “extrema” in (Stine et al., 2020)), subjects respond to the first brief burst of strong evidence they detect (Figure 3a, see also (Cartwright and Festinger, 1943; Ditterich, 2006; Hyafil et al., 2023)). This is modeled by continuously integrating all clicks within a short window. If all the clicks within the window favor the same response and the number of clicks hits or surpasses a threshold, the model commits to that response and ignores all subsequent clicks (eliminating the requirement that all clicks in the window favor the same response does not alter any of the findings discussed below). If no sufficiently strong burst of clicks is detected by the end of the stimulus, the model produces a 50/50 guess (Figure 3a example trials 1, 6, and 7). In principle the bust strategy is capable of resulting in response behavior consistent with the analysis in Figure 2, thereby mimicking full integration. For example, trials with a greater overall click difference are more likely to contain a burst event favoring the correct response, resulting in a monotonically increasing psychometric plot. Trials with longer stimuli are also more likely to contain a burst event, resulting in fewer guesses and an increasing chronometric plot. Finally, if on average burst events occur at a random time once per trial, the period within the stimulus sampled will be uniform leading to a flat psychophysical kernel. Therefore, if these assumptions are true, a cursory analysis of behavioral data appears insufficient to rule out if a subject is using the burst strategy. Below we will perform additional analysis to test if rats are using the burst strategy when performing the Poisson Clicks Task.

**Figure 3.**
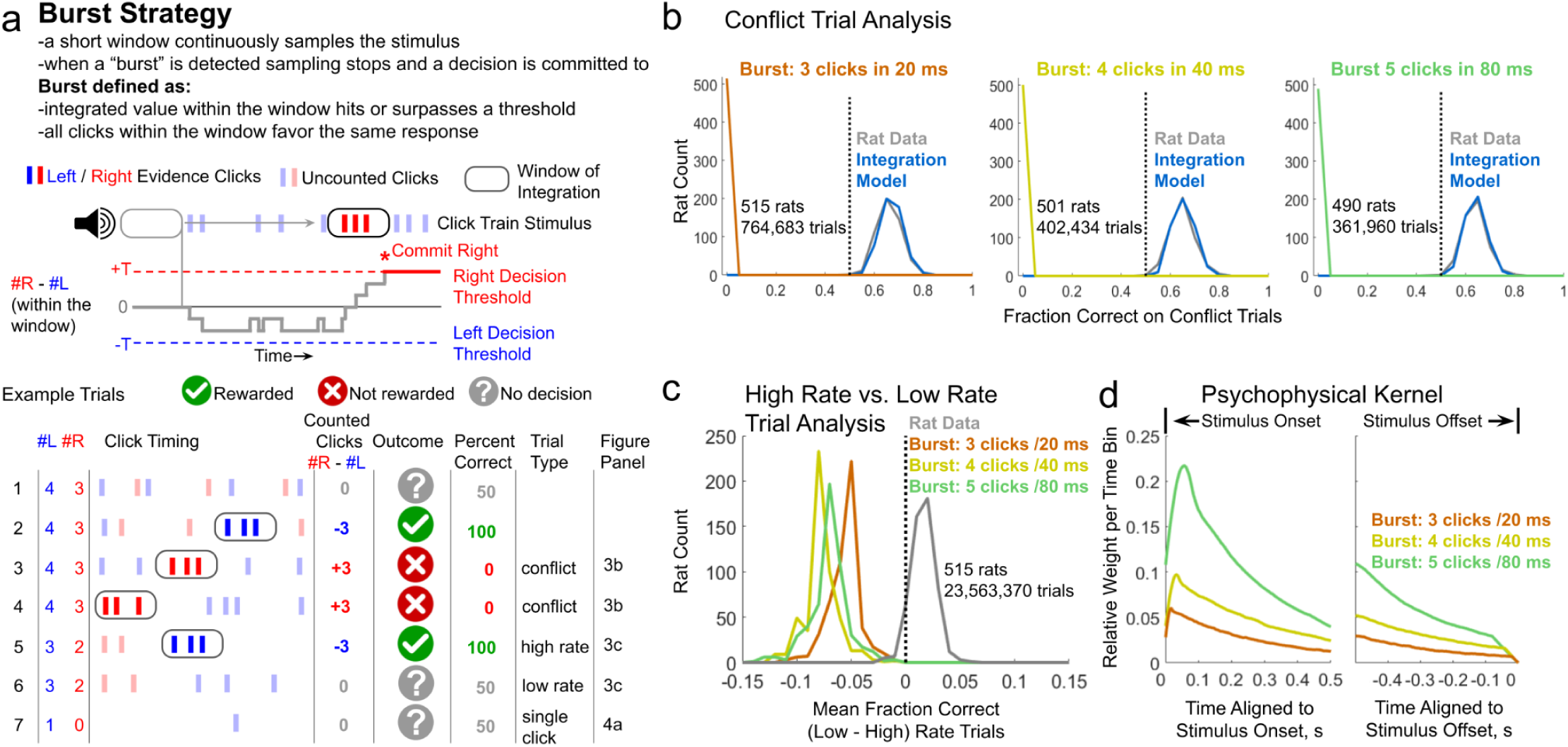
Analysis of performance of the Burst Strategy on the Poisson Clicks task. **a)** Upper: Schematic describing the burst strategy. A short window samples the stimulus. When all the clicks within the window favor the same response and their total hits or surpasses a threshold, the model commits to a decision and ignores all subsequent clicks. Lower: Example trials demonstrating how the burst strategy performs given different stimuli. In these examples the threshold is 3 clicks within the window of integration. Trials where the model commits to no decision result in a 50/50 guess. The percent correct indicates how this threshold and window duration perform on these example trials. The trial types refer to later panels in this and subsequent figures, **b)** Conflict trials are those a strategy is defined to get wrong more than 50% of the time. Here we test three variants of the burst model: Left panel, 3 clicks in 20 ms; Center panel, 4 clicks in 40 ms; Right panel, 5 clicks in 80 ms. In the identified conflict trials, the first “burst” of clicks is from the “incorrect” side and therefore the strategy will always get these trials wrong (see (a) example trials 3 and 4). Histogram of the mean performance on the identified conflict trials plotted across rats for the burst strategy, integration model, and the rats’ data. Rats and the integration model perform better than chance on the conflict trials, consistent with them not using the burst strategy to perform the task. Only rats with at least 20 conflict trials are included in this analysis, **c)** Analysis of rat and model performance on trials with higher and lower overall click rates. First, trials are divided into groups with each particular # left and # right clicks, i.e. one group will contain all the trials with 8 left and 5 right clicks, while another group contains all the trials with 10 left and 15 right clicks. Second, each of those groups is split based on if the stimulus duration is greater or less than the group’s median. Shorter duration trials are flagged as “high rate trials” and longer duration trials are flagged as “low rate trials” (see **(a)** example trials 5 and 6). Histogram is plotted across rats of the difference in fraction correct on low versus high rate trials. In general rats perform slightly better on low rate trials. The burst strategy predicts the opposite, better performance on high rate trials. This is because high rate trials are more likely to contain a burst, thereby leading to fewer guesses, **d)** Psychophysical kernels generated from 3 variants of the burst model run on trials at minimum 500 ms long generated with the same statistics as the trials the rats performed. Trials are aligned to either the stimulus onset, left plot, or offset, right plot. All variants result in a decaying kernel since any burst detected early in a trial prevents all subsequent clicks from being sampled. The brief rise during the earliest part of the stimulus period is a result of the duration of the window of integration sampling clicks over a finite period of time, similar to the boundary effects seen in sampling from the snapshot strategy (see Figure 5d).

In the first analysis we look at model and rat performance on a subset of trials we refer to as “conflict trials”. Conflict trials are named because the presented evidence conflicts with the model’s response. The particular timing of the clicks in these trials is such that the degenerate strategy is more likely to sample the incorrect clicks and get the trial wrong (Figure 3a example trials 3 and 4). For example, Figure 3a example trial 3, while there are more left clicks, the fewer right clicks occur in a burst and trigger an incorrect decision and response. In example trial 4, again with more left clicks, both left and right clicks occur in bursts but the right burst happens first, again triggering an incorrect response. Both of these are examples of conflict trials. In figure 3b we test 3 variants of the burst model with different duration windows and burst thresholds: 3 clicks in 20 ms, 4 clicks in 40 ms, and 5 clicks in 80 ms (we tested all combinations of window of integration duration 10, 20, 40, and 80 ms, with burst threshold 3, 4, 5, and 6 clicks and in all cases found similar results to the three variants shown here). Given the burst threshold we identify conflict trials and plot a histogram across rats of their mean performance on these trials along with the performance of the burst strategy and a model that integrates the entire stimulus. Only rats that have a minimum of 20 conflict trials are included in this analysis. By design the burst strategy always gets conflict trials wrong (Figure 3b), as they are selected to have the first burst of clicks favor the incorrect response. If rats also performed below chance on these trials, that would be evidence they are responding to these burst events. However, in all 3 variants of the burst strategy tested all rats perform above chance on these trials, consistent with them not responding to the bursts of clicks. In contrast, the integration model is able to match the rats’ performance on these trials.

For a second test we exploit the inherent noise in the timing of individual clicks (similar to analysis performed in (Brunton et al., 2013)). While the total generative click rate (left rate + right rate) is typically 40 Hz (some rats performed 20 Hz trials), because of the Poisson nature of the click timing, some trials will by chance have a total rate greater than 40 Hz while other trials it will be less. Here we divide the trials into two equal groups differing only in the overall rate. First we divide the trials into groups with a particular number of left and right clicks, i.e. all 3 left / 5 right click trials go into one group while all 7 left / 4 right click trials go into another. Each group is then divided in two based on the stimulus duration being greater than or less than the median for that group. The “short duration” trials go into the “high rate” group and the “long duration” trials go into the “low rate” group. Four types of trials are excluded from this analysis: groups with only 1 trial; groups with an equal number of left and right clicks; in groups with an odd number of trials, the trial with stimulus duration equal to the group median is excluded; and all frozen noise trials are excluded. This results in two groups of trials with an equal representation of all particular click counts while differing in the total click rate. Longer duration trials have an overall lower total click rate, are less likely to contain a burst event, and therefore more likely to require a 50/50 guess at the end of the stimulus compared with shorter duration (high click rate) trials. Therefore, a burst strategy predicts lower performance on low rate trials compared to high rate trials. Analysis of the 3 variants of the burst model tested against the trials presented to the rats confirm this prediction (Figure 3c). However, analysis of the rats’ performance in these two groups of trials actually shows a slightly better performance on low rate trials compared to high rate trials, a finding inconsistent with rats using a burst strategy to perform the Poisson Clicks Task.

The third test we reexamine the premise that a burst strategy can uniformly sample the stimulus and result in a flat psychophysical kernel matching the mean of the rats’ data seen in Figure 2c. This requires that each trial contains no more than one burst and that burst must be randomly positioned within the stimulus. Given the Poisson nature of click times used to generate the stimuli in this task, that premise is not possible. Any event that occurs on average once per trial will follow a Poisson distribution with lambda = 1. Therefore, we’d expect 36.8% of the trials to have no bursts, and 26.4% of trials to have more than one burst event. Since the model commits to a decision at the first burst event it encounters, on trials with more than one burst the later bursts will be ignored, leading to an over sampling of earlier clicks in the stimulus. To achieve more uniform sampling requires a lower lambda such that few trials have more than one burst event. A Poisson distribution with lambda = 0.355 results in only 5% of trials having more than one burst. However, such a low lambda corresponds to a relatively rare burst event. Now ∼70% of the trials will have no burst event, trials on which a subject using the burst strategy must guess. The best performance one can achieve in this regime, assuming all trials with a burst the burst favors the correct side, which they do not, is ∼65%; yet 65% correct was the minimum criteria for rats to be included in this data set, and on average rats in this dataset perform the Poisson Clicks Task much better averaging 76.2% correct (95% perform between 68.3% and 83.3% correct).

To demonstrate the magnitude of oversampling of early clicks the burst strategy results in, in Figure 3d we analytically compute the psychophysical kernel that would result from the 3 variants of the burst model described previously. The models were run on simulated trials generated with the same statistics as the trials the rats typically performed and with a minimum duration of 500 ms. Here we see a robust decrease in sampling over time, as expected since any early burst prevents all later clicks from being sampled. This is not a consequence of the particular parameters chosen for these model variants but a necessary consequence of the Poisson nature of the click timing. The rise in sampling probability at the earliest time points is a consequence of the finite duration of the window of integration (similar to what is seen later in the snapshot strategy Figure 5d).

Unlike continuous evidence integration tasks, the Poisson Clicks Task contains discrete packets of information, with some trials containing as little as one non-stereo click. For the fourth test we examine trials that only contain a single click (ignoring the stereo click that begins some stimuli). While the criteria for what constitutes a “burst” can be debated, a single click would generally not meet that definition. Therefore if a subject was using the burst strategy to perform the Poisson Clicks Task one would predict they would guess on these trials. Each of the 3 variants of the burst model discussed above would never hit threshold on a single click trial and therefore we predict if a subject was using a burst strategy to perform the Poisson Clicks Task they should respond at chance on all single click trials. Rats however are able to perform significantly above chance on these trials (∼65% correct, Figure 4a, only rats that were presented at least 20 single click trials are included in this analysis). In order to match the rats’ performance on single click trials, noise needs to be added to the model such that a single click surpasses the burst threshold ∼30% of the time (30% of trials the subject gets correct, 70% they guess yielding 35% correct, for an overall correct of 65%). However, if the model commits to a decision 30% of the time on single click trials, the model will commit to a decision 30% of the time it encounters any click on all other trials. This would lead to an even steeper decaying psychophysical kernel from what was shown in Figure 3d. There is no way to reconcile these two features of rats’ performance on the Poisson Clicks Task, above chance performance on single click trials and a flat psychophysical kernel, if they are using only a burst strategy.

**Figure 4.**
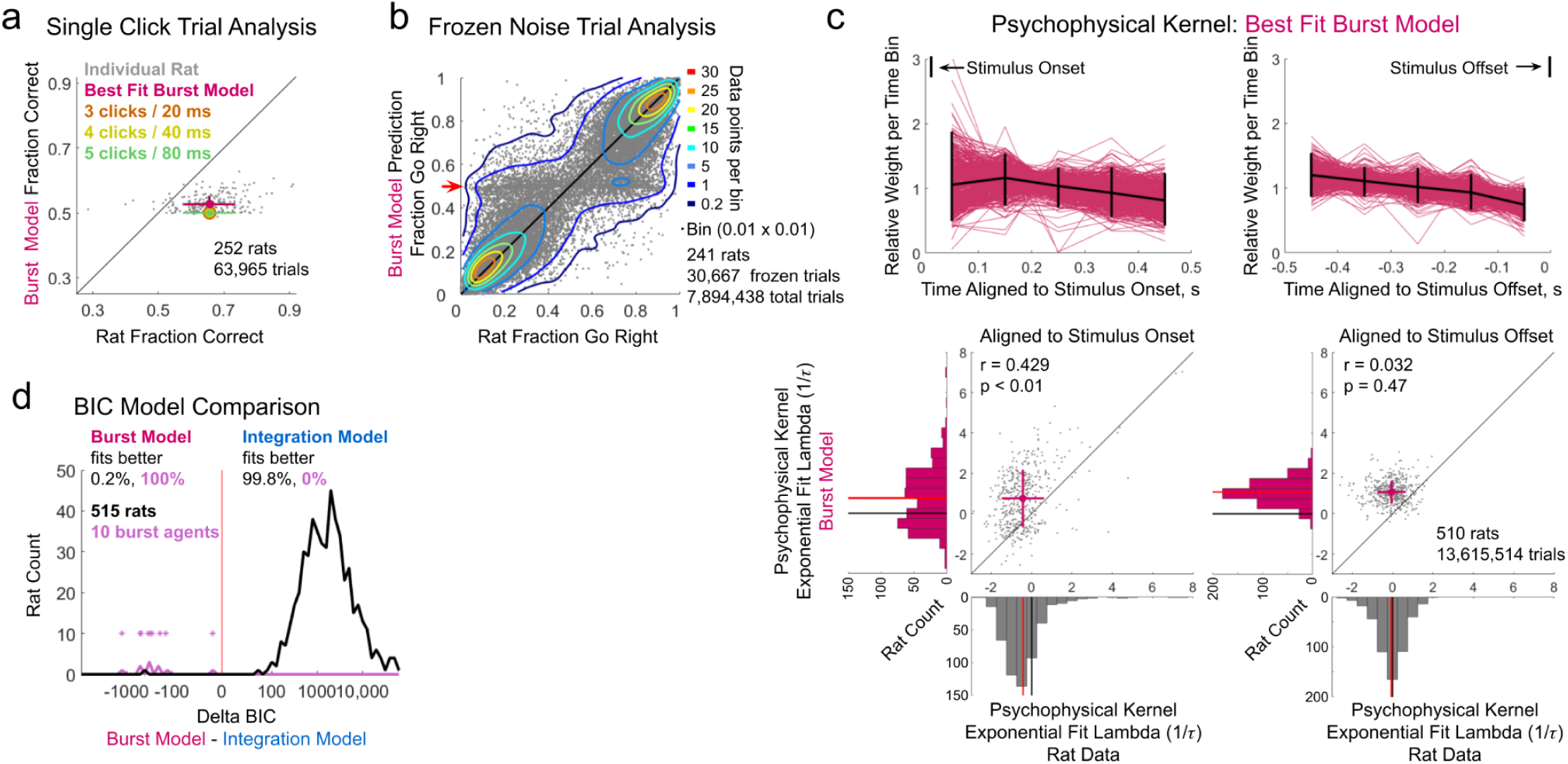
Analysis of the best fit Burst Model to rat performance on the Poisson Clicks task. **a)** Rat and burst model performance on trials with only a single click (ignoring the stereo click that begins some trials). Individual dots represent mean performance from individual rats that experienced at least 20 single click trials versus performance of the best fit burst model. Mean performance of three variants of the burst model shown in color (the three data points overlap), **b)** Frozen noise trials are repeated trials with identical click train stimuli. The burst model is fit to all non-frozen trials, i.e. those that have unique stimuli, and tested on frozen trials. Not all rats experienced frozen noise trials when performing the Poisson Clicks task. Only rats that were presented at least 40 different frozen noise trials, each at least 100 times, are included in this analysis. Data points represent an individual rat versus model predicted performance across a single unique frozen noise trial set. Scatter plot density is indicated with contour lines. Bins are 0.01 x 0.01 fraction go right. Red arrow indicates band of trials the best fit burst model predicts near 50% performance but rats produce a large range of responses further from 50%. c) Top: The psychophysical kernel produced by the best fit burst model to each individual rat (red) aligned to stimulus onset (left) or offset (right) for trials with stimuli longer than 500 ms. Mean shown in black. Vertical bars indicate 95% of the rats. Bottom: An exponential function is fit to each psychophysical kernel produced by the best fit burst model for each rat. Plot of the lambda value (1/r) from this exponential fit compared to the lambda of the fit to each rat’s psychophysical kernel (Figure 2c). Left aligned to stimulus onset, right aligned to stimulus offset. Histograms of the lambda values plotted separately for the burst model and rats’ data. Rat data histograms are the same as in Figure 2c. The burst model results in psychophysical kernels that are generally decaying (lambda > 0) significantly more so than the rats’ psychophysical kernels (aligned to onset p < 0.01; aligned to offset p « 0.01). The duration of the window of integration can result in a rising kernel at the earliest time points lowering the lambda value when aligned to stimulus onset (see Figure 3d), **d)** Histogram of delta BIC comparing the full integration model (10 parameters) with the full burst model (9 parameters). See Table 1 for a list of parameters included in each model. Only 1 rat (0.2%) is better fit by the burst model. Synthetic choice data was generated using a range of parameters on the burst strategy. All 10 of these synthetic agents (10k trials each) were better fit by the burst model. Individual synthetic data

For the final set of tests of the burst strategy we fit an extended version of the model, containing the same free parameters as the integration model, to the rats’ data (see Table 1 for a list of parameters, since the burst strategy does not integrate over long periods of time we eliminated 2 parameters, *σ*_a_ and *λ*, and the bias term was replaced by the burst threshold). Even after fitting a model to the rat’s choice data across all trials we see the best fit burst strategy is still not able to match rats’ performance on single click trials (Figure 4a).

**Table 1.** Parameters used in model variants. Models (columns) can have up to 11 free parameters (rows). Parameters that are fit by the model are indicated with “free”. Parameters that have a fixed value are indicated with that value. Parameters that are excluded altogether from a model are indicated with an “X”. The bottom row indicates the figures in which the model is used. The rightmost column is a brief description of what each parameter does (see Methods for complete description).

| Model | Burst | Burst Extended | Snapshot | Snapshot Extended | Single Click | Single Click Extended | Integration | Parameter Description |
| --- | --- | --- | --- | --- | --- | --- | --- | --- |
| Parameter |  |  |  |  |  |  |  |  |
| $\sigma_i$ | X | free | X | free | X | free | free | standard deviation of the distribution of the accumulator’s initial value, in clicks |
| $\sigma_s$ | X | free | X | free | X | free | free | Standard deviation of the value of a single click, in clicks |
| $\sigma_a$ | X | 0 | X | free | X | X | free | accumulator diffusion, standard deviation after 1 second, in clicks |
| Bias (burst threshold) | 3, 4, 5 | free | X | free | X | free | free | Decision criteria bias (burst model threshold), in clicks |
| Lapse | X | free | X | free | X | free | free | Fraction of trials the model ignores the stimulus |
| Lapse Bias | X | free | X | free | X | free | free | Fraction of right responses on lapse trials where the model ignores the stimulus |
| $\alpha$ | X | free | X | free | X | X | free | Controls the exponent the accumulated value of clicks is raised to |
| $\phi$ | X | free | X | free | X | X | free | Click adaptation, how much one click affects the magnitude of subsequent clicks |
| $\tau_\phi$ | X | free | X | free | X | X | free | Time constant of click adaptation |
| Window of Integration | 20, 40, 80 | free | 100, 250, 500 | free | 50, 125, 250 | free | 1000 | Duration of a window the model integrates clicks within, in milliseconds |
| $\lambda$ | X | 0 | X | free | X | X | free | Inverse of the accumulator’s memory time constant |
| Figures Used In | 3b,c,d<br>4a | 4a-d | 5b,c,d<br>6a | 6a-d | 7b,c,d<br>8a | 8a-d | 4d, 6d, 8c,d<br>9, 10, 11 |  |

In the fifth test we fit the extended burst model to the rats’ performance on only the unique (non-frozen) trials, then tested the model’s predicted response against the rats’ choice behavior on the frozen noise trials (Figure 4b). Here we only include rats that had been presented at least 40 different frozen trials and only frozen trials that had been presented at least 100 times each to the rat. In total 241 rats and 30,667 different frozen trials met this criteria. Each of these trials were presented on average 257 times to each rat (therefore this plot contains data from 7,894,438 total trials). Since the scatter plot in Figure 4b contains over 30,000 points we overlaid contour lines to allow one to assess the density of data points. While there is an overall correlation between the rats’ response behavior and the model’s prediction, there is also a large cloud of points centered on the model predicting ∼50% respond “right” but spread across the full range of rat responses (indicated with a red arrow along the vertical axis). These are trials where the model does not detect a burst and must guess, but where the rats are consistently producing responses further from 50%.

On average the correlation between the rat’s performance and the model’s predicted responses is r = 0.927 with the median difference between the predicted and actual performance being +/- 5.8%. A small fraction of the frozen noise trials are poorly fit by the model as indicated by the data points that lie far from the diagonal (6.7% of trials have a difference between the model’s predicted response and the rats’ actual response greater than 25%). For comparison the integration model discussed later (Figure 9e) more accurately predicts choices on these trials (correlation r = 0.951, median difference between predicted and actual performance 4.4%, and 3.0% of trials have a difference between the model’s predicted and the rats’ actual response > 25%).

In the variants of the burst strategy discussed earlier, we argued that it is not possible for the burst strategy to result in a flat psychophysical kernel given the Poisson nature of the click timing. However, the 3 variants tested in Figure 3 did not have all the free parameters in the extended burst model. In Figure 4c top we plot the psychophysical kernels generated by the best fit burst model aligned to stimulus onset (left) and offset (right). As in Figure 2c we again fit an exponential function to each kernel and plot the lambda value (1/*τ*) from these fits compared to the lambda values from the rats’ actual psychophysical kernels (Figure 4c bottom). Regardless whether the trials are aligned to stimulus onset or offset, the burst model results in kernels that have a greater lambda value (are more decaying) than the kernels fit to the rats’ actual response behavior (p < 0.01 for both alignments). This effect, while still significant, is less pronounced when trials are aligned to stimulus onset. This is because longer windows of integration can result in a briefly rising probability of sampling clicks at the earliest time periods (see Figure 3d) that then decays for the remainder of the stimulus period.

Finally, for our seventh test of the burst strategy we use Bayesian Information Criterion (BIC) analysis (Schwarz, 1978) to directly compare how well the burst and integration models fit the rats data (Figure 4d, black, see Methods). Here we find that 99.8% (514 out of 515) of the rats are better fit by the integration model when compared to the extended burst model. As a test of the model fitting and this analysis we generated 10 synthetic agents and simulated their responses using the burst strategy (Palminteri et al., 2017; Wilson and Collins, 2019). Each agent used different parameters run through the extended burst model on the first 10,000 trials that were presented to rat K209 (this was the minimum number of trials a rat must have completed to be included in this dataset). All 10 of the synthetic agents were better fit by the burst model (Figure 4d, purple), consistent with this analysis being able to identify which strategy is more in line with what is being used by the rats.

In total, rats’ behavior is inconsistent with the burst strategy. First, the rats do not perform below chance on conflict trials (Figure 3b), trials where the first burst of clicks favors the incorrect response. Second, they do not show the decrease in performance on low rate trials compared to high rate trials (Figure 3c) as expected by the model. Third, the rats do not show a steeply decaying psychophysical kernel (compare Figures 2c and 3d) as required by a burst strategy used to perform an accumulation of evidence task where the stimuli are generated using Poisson statistics. Fourth, rats perform significantly above chance on trials with only a single click (Figure 4a), trials that contain no bursts. Fifth, the rats produce consistent responses on a subset of frozen noise trials that the burst model is only able to guess on (Figure 4b red arrow). Sixth, even the best fit extended burst model results in psychophysical kernels that are significantly more “decaying” that those fit to the rats’ response behavior (Figure 4c). Seventh, BIC analysis favors the rats using full integration over a burst strategy. Taken together, these analyses demonstrate that not only is it possible to determine if a subject is using a burst strategy to solve the Poisson Clicks Task, but that overall the rats in this dataset are not.

### Testing the Snapshot Strategy

The snapshot strategy supposes that rather than integrating the entire stimulus a subject only integrates a randomly selected short portion of the stimulus to inform their decision (Figure 5a) (Hyafil et al., 2023; Pinto et al., 2018; Stine et al., 2020). This is modeled by placing a short window of integration randomly within the stimulus period; all clicks within that window are integrated, while any clicks outside the window are ignored. If no clicks are present within the window the model simply produces a guess (Figure 5a, example trials 5 and 7). In principle the snapshot strategy is capable of producing response behavior consistent with the analysis in Figure 2, thereby mimicking full integration. The greater the click difference the more likely the snapshot will sample more “correct” clicks, resulting in a monotonically increasing psychometric plot. Since the duration of any given stimulus is not known *a priori*, the snapshot may end up being placed over silence prior to stimulus onset on shorter duration trials. Therefore the longer the stimulus, the more likely the snapshot will overlap clicks, resulting in increased performance and a rising chronometric plot. Finally, if the snapshot is brief enough it can uniformly sample the entire stimulus period leading to a flat psychophysical kernel. As with the burst strategy, if these assumptions are true, a cursory analysis of behavioral data appears insufficient to determine if a subject is actually integrating the entire stimulus. Below we will perform a similar set of analyses used above for the burst strategy to determine if rats are using a snapshot strategy when performing the Poisson Clicks Task.

**Figure 5.**
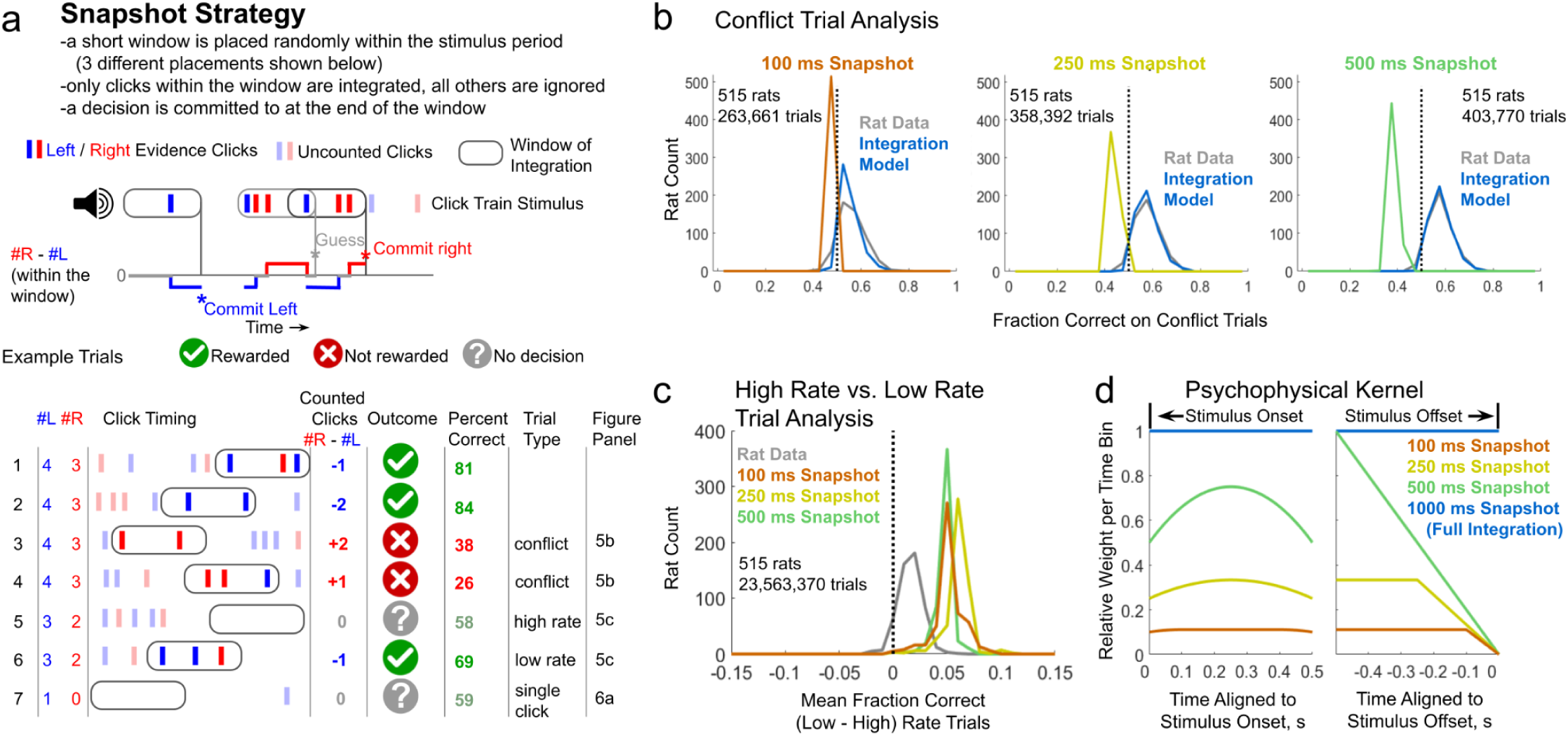
Analysis of performance of the Snapshot Strategy on the Poisson Clicks task. **a)** Upper: Schematic describing the snapshot strategy. A short window is placed randomly within the stimulus period (here 3 possible placements are shown which lead to different outcomes). All clicks within the window are integrate and a choice is committed to. All clicks outside the window are ignored. Lower: Example trials demonstrating how the snapshot strategy performs given different stimuli. Trials where the model samples no clicks result in a 50/50 guess. The outcome indicates how the snapshot strategy performs given the particular window placement shown while the percent correct indicates the mean performance of the snapshot strategy across all possible window placements on these example trials. The trial types refer to later panels in this and subsequent figures, **b)** Conflict trials are those the snapshot strategy is defined to get wrong. Here we test three variants of the snapshot model: 100, 250, and 500 ms windows of integration. Conflict trials are those with particular click times such that the window of integration is more likely to sample incorrect clicks (see (a) example trials 3 and 4). Histogram of the mean performance on the identified conflict trials plotted across rats for the snapshot strategy, integration model, and the rats’ data. Rats and the integration model perform better than chance on the conflict trials, consistent with them not using the snapshot strategy to perform the task. Only rats with at least 20 conflict trials are included in this analysis, c) Analysis of rat and model performance on trials with higher and lower overall click rates. First, trials are divided into groups with each particular # left and # right clicks. Second, each of those groups is split based on if the stimulus duration is greater or less than the group’s median. Shorter duration trials are flagged as “high rate trials” and longer duration trials are flagged as “low rate trials” (see (a) example trials 5 and 6). Histogram is plotted across rats of the difference in fraction correct on low versus high rate trials. In general rats perform slightly better on low rate trials. The snapshot strategy predicts a larger difference in performance since low rate (long) trials have less silence before the stimulus starts and therefore there’s less chance the window will sample silence leading the model to guess, d) Psychophysical kernels computed for 3 variants of the snapshot model run on trials at minimum 500 ms long generated with the same statistics as the trials rats performed. Trials are aligned to either the stimulus onset, left plot, or offset, right plot. Only the very long or very short windows of integration approach a relatively uniform sampling of the stimulus period. All except full integration under sample the earliest and latest portions of the stimulus period.

As a first test of the snapshot strategy, we identify “conflict” trials, trials with particular click timing such that the window of integration is more likely to sample incorrect clicks and therefore get the trial wrong (Figure 5a, example trials 3 and 4). Unlike with the burst strategy, here the snapshot strategy does not always get conflict trials wrong, it’s simply more likely to do so. This is because of two reasons: 1) clicks that are clustered together occupy less temporal space and therefore are less likely to be sampled by a random snapshot; 2) clicks occurring at the beginning and end of the stimulus period are less likely to be sampled given moderate length snapshots (see Figure 5d). So, even though these trials have more left clicks, since those clicks tend to either cluster together or occur nearer the beginning and end of the stimulus period, or both, the model is more likely to sample right clicks and respond “right” getting the trial wrong. In Figure 5b we identify conflict trials for three variants of the snapshot model using a window of integration of 100, 250, or 500 ms in duration and test how the snapshot strategy performs on these trials compared with the rats and an integration model. The snapshot models by design perform below chance on the conflict trials, as this is how the trials were selected. However, when we look at the rats’ performance on these trials we see they perform significantly above chance, as does the integration model. The particular timing of the clicks in these trials are such that if a subject was using a snapshot strategy they should get these trials wrong. Since the vast majority of rats perform above chance on these trials (Figure 5b, 90.3%, 96.5%, and 96.7% of rats perform above chance on 100, 250, and 500 ms snapshot conflict trials respectively), that’s evidence they are not using a snapshot strategy to perform the Poisson Clicks Task.

In a second test we compare the performance of the rats and models on trials with overall higher or lower total click rates. As previously, we divide the trials into two groups, where each group contains an equal representation of trials with each particular number of right and left clicks, but differ in the rate at which those clicks are presented (see burst strategy section above for more details). In general the snapshot strategy predicts better performance on low rate compared to high rate trials (Figure 5c). This is because the window of integration has a greater chance of overlapping clicks rather than silence the more spread out those clicks are. Rats do show a small improvement in performance on low versus high rate trials, but this is significantly smaller than the difference predicted by the snapshot strategy.

When we introduced the snapshot strategy we stated that if the snapshot is brief enough it can uniformly sample the entire stimulus period. However, because the stimulus in the Poisson Clicks Task is delivered at discrete times, extremely brief snapshots are increasingly likely to only sample the silence between clicks. Even after ignoring the silence before the stimulus onset, a 10 ms window of integration has a ∼64% chance of overlapping no clicks given a 40 Hz total generative rate. Therefore, longer windows of integration are required to ensure that clicks are sampled. In Figure 5d we plot the analytically computed psychophysical kernels for the variants of the snapshot model discussed so far when run on trials with a minimum duration of 500 ms. While very short windows of integration are able to sample the stimulus period relatively uniformly, longer duration windows result in boundary effects undersampling the earliest and latest portions of the stimulus period. This is best appreciated if one imagines a 500 ms snapshot though it holds for all moderate duration snapshots. Given all possible placements of a 500 ms window of integration within a 1s stimulus period, any clicks that occur near the middle will always be sampled since all placements of the window of integration overlap the middle of the stimulus, while those nearer the beginning and end less so since few placements of the window overlap these periods. The psychophysical kernels fit to the rats’ data (Figure 2c) show no consistent underweighting of the early and late portion of the stimulus period. Therefore the rats’ flat psychophysical kernels are inconsistent with them using the snapshot strategy with a moderate duration window of integration.

One possible means to eliminate the boundary effects seen when moderate duration windows of integration are confined to the stimulus period is to allow the window to extend beyond those bounds. Allowing the window to begin prior to the start of the stimulus period or terminate after the end of the stimulus period by a duration equal to the length of the window of integration would produce perfectly uniform sampling within the stimulus period for any duration window of integration. However, apart from the questions this strategy raises; Why would one place a window of integration when they know there will never be stimuli?; it also has the consequence of lowering the probability of sampling clicks at any given time. While a 500 ms window of integration confined to the stimulus period will sample each period within the stimulus 50% of the time, though not uniformly, the same duration window now free to extend beyond the stimulus period will sample each period within the stimulus only 33% of the time. When the window of integration is confined to be within the stimulus period, as it is extended, the snapshot strategy approaches full integration, and the two models are equivalent when the snapshot is 1000 ms long. However, if the window is allowed to extend beyond the stimulus period, even a 1000 ms snapshot will only sample each portion of the stimulus 50% of the time. An infinite duration window of integration would be required to be equivalent to full integration. This will have consequences for the analysis of single click trials discussed below (Figure 6a).

**Figure 6.**
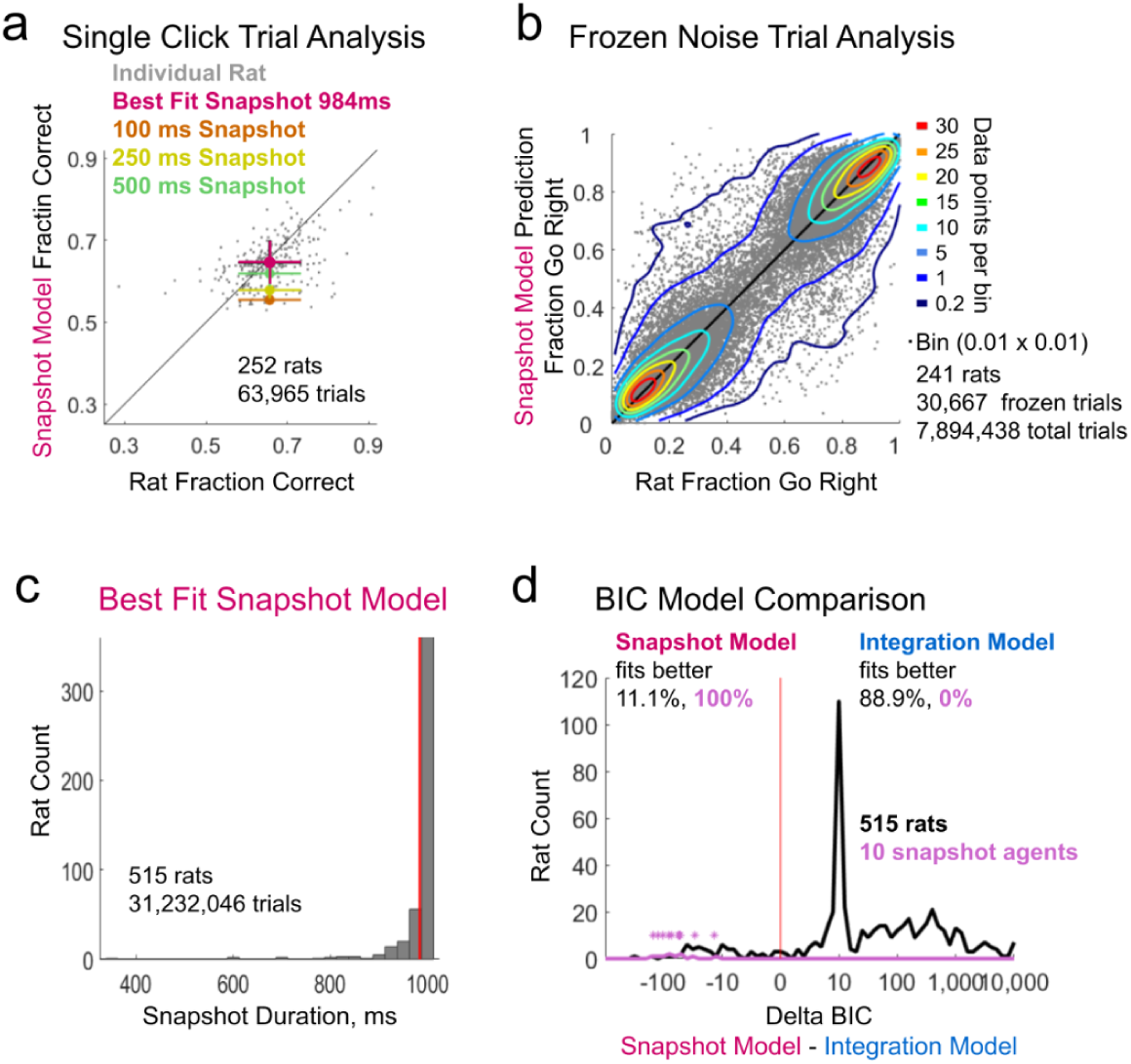
Analysis of the best fit Snapshot Model to performance on the Poisson Clicks task. **a)** Rat and snapshot model performance on trials with only a single click (ignoring the stereo click that begins some trials). Individual dots represent individual rats that experienced at least 20 single click trials versus the best fit snapshot model. Mean performance of three variants of the snapshot model shown in color, b) Frozen noise trials are repeated trials with identical click train stimuli. The snapshot model is fit to all non-frozen trials, i.e. those that have unique stimuli, and tested on frozen trials. Data points represent an individual rat versus model predicted performance across a single unique frozen noise trial set. Scatter plot density is indicated with contour lines. Bins are 0.01 x 0.01 fraction go right, **c)** Histogram of the best fit value of the window of integration duration parameter when the full snapshot model is fit to the rats’ data. Mean of the distribution plotted in red. The best fit snapshot model approaches full integration with a mean duration for the window of integration at 984 ms (full integration = 1000 ms) d) Histogram of delta BIC analysis comparing the full integration model (10 parameters) with the full snapshot model (11 parameters). See Table 1 for a list of parameters included in each model. Rats with a negative delta BIC (11.1%) are better fit by the snapshot model. However, on average the best fit window of integration is 923 ms for these individuals. Synthetic choice data was generated using a range of parameter values run on the snapshot strategy (window of integration duration ranged from 50 to 500 ms). All 10 of these synthetic agents (10k trials each) were better fit by the snapshot model. Individual synthetic data points also indicated with * as their histogram can be obscured.

As with the burst strategy tested above, for this final set of tests of the snapshot strategy we fit an extended version of the model to the rats’ data (Table 1). The extended version of the snapshot model contains the same free parameters as the integration model with the addition of a parameter controlling the duration of the window of integration. This parameter is free to adopt any value between 1 and 1000 ms. As it approaches 1000 ms the snapshot model converges to become the full integration model.

The snapshot strategy is able to replicate performance with a rising chronometric plot (Figure 2b) because longer duration trials have a shorter period of silence preceding the stimulus and therefore there’s a smaller period where the snapshot may sample no clicks. The extreme extension of this are trials with only a single click. These trials have the largest period of silence and therefore the shorter the duration of the window of integration the closer to chance performance should be on single click trials. In Figure 6a we show the mean performance on single click trials expected if rats were using snapshots of varying durations. Only as one approaches a 500 ms window of integration does performance of the snapshot strategy begin to match the rats’ performance. Given a total click rate of 40 Hz, a 500 ms window would contain upwards of 20 clicks, while this is not integrating the entire stimulus, it is still a significant amount of integration which is at odds with the premise of a model taking a “snapshot” of the stimulus (Stine et al., 2020). Allowing the window of integration to extend beyond the stimulus period as discussed above would lower the probability of sampling any given period within the stimulus, and therefore would require an even longer window to match rats’ performance. The 246 rats that experienced at least 20 single click trials, performed on average ∼65% correct on them. The extended snapshot model is able to match this performance well (Figure 6a) but only because the best fit duration for the window of integration is so high (∼984 ms, see Figure 6c).

Next we repeated the test using frozen noise trials, now comparing the predicted performance of the snapshot model against the rats’ responses. Briefly, the extended snapshot model was fit to the rats’ performance on only the unique (non-frozen) trials. We then used the best fit parameters to predict performance on the repeated frozen noise trials and plotted that against the rats’ actual performance (Figure 6b). On average the extended snapshot model is able to predict the rats’ actual response behavior well, however, given that it has one additional free parameter when compared with the integration model, and is able to converge to the integration model as the duration for the window of integration increases, the snapshot model actually predicts performance on frozen noise trials slightly worse than the integration model. On average the correlation between the rat’s performance and the model’s predicted responses is r = 0.950, the median difference between the predicted and actual performance being +/- 4.5%, and 3.2% of trials are poorly predicted having a greater than 25% difference between the predicted and actual response. For comparison the integration model with one less free parameter (discussed later see Figure 9e) has a correlation r = 0.951, median difference between predicted and actual performance of 4.4%, and 3.0% of trials are poorly predicted.

In Figure 6c we plot a histogram of the duration of the window of integration from the best fit extended snapshot model across rats. The average duration across rats is 984 ms (mean) and 998 ms (median), with 96% of rats being best fit with a snapshot duration greater than 900 ms and only one rat less than 500 ms. While still technically following the rules of the snapshot strategy, such a long window of integration runs counter to its premise, integrating over 900 ms of an at most 1000 ms stimulus is not responding to a “snapshot” of the stimulus.

Finally we directly compare the goodness of fit between the snapshot and integration models to the rats response data (Figure 6d, black) using BIC analysis. We find that 88.9% of rats are better fit by the integration model. Of the 11.1% of rats better fit by the snapshot model, they are best fit with a mean snapshot duration of 923 ms. As done previously when testing the burst strategy, here we generated 10 synthetic snapshot agents, using a range of snapshot durations, and simulated their choices on the same 10,000 trials. When comparing the integration and snapshot model fits to these synthetic datasets we find the BIC analysis favors the snapshot model over integration for all of them (Figure 6d, purple), validating the model fitting and this analysis.

Taken together, rats’ behavior is inconsistent with them using a snapshot strategy. First, the rats do not perform below chance on conflict trials (Figure 5b), trials that contain particular click timing such that a snapshot is more likely to sample incorrect clicks. Second, they do not show the increase in performance on low rate trials compared to high rate trials of the magnitude expected by the model (Figure 5c). Third, the rats do not show consistent symmetric edge effects in the psychophysical kernels undersampling both the earliest and latest periods within the stimulus (compare Figures 2c and 5d). Fourth, they perform significantly above chance on trials with only a single click (Figure 6a), necessitating at a minimum a window of integration at least 500 ms in duration. Fifth, the snapshot model, despite being the integration model plus one additional free parameter, predicts performance on frozen noise trials slightly worse than the integration model itself (Figure 6b). Sixth, the best fit extended snapshot model adopts a strategy nearly identical to integration with a mean window of integration of 984 ms for stimuli that are at a maximum 1000 ms in duration (Figure 6c), inconsistent with the premise of the model. Seventh, BIC analysis favors the vast majority of rats using full integration over a snapshot strategy. These analyses demonstrate that not only is it possible to determine if a subject is using a snapshot strategy to solve the Poisson Clicks Task, but that overall the rats in this dataset are not.

### Testing the Single Click Strategy

The single click strategy combines elements of the snapshot and burst strategies discussed above. As with the snapshot strategy, a window of integration randomly begins within the stimulus period. However, rather than integrating all clicks within that window, only the first click is considered, equivalent to a burst threshold of one (Figure 7a). If no clicks occur within the window the model guesses (Figure 7a, example trial 5). The window of integration ends when either a click is detected, the duration for the window of integration is reached, or the stimulus period ends. While only considering a single click on each trial, this strategy is still in principle able to replicate the 3 behavioral analysis from Figure 2. The greater the click difference the more likely the single click attended to will be “correct”. This will result in a monotonically increasing psychometric plot. As with the snapshot strategy, the longer the stimulus duration, the greater the chance the window of integration will sample a click rather than silence, resulting in an increasing chronometric plot. Finally, since clicks have a relatively uniform probability of being sampled regardless of when they occur during the stimulus period, the single click strategy should result in a flat psychophysical kernel. As with the other degenerate strategies considered here, if these assumptions are true, simply looking at these three plots (psychometric, chronometric, and psychophysical kernels) is not enough to determine which strategy rats are using to solve the Poisson Clicks Task. Below we will perform the same analyses used previously for both the burst and snapshot strategies to determine if rats are using the single click strategy when performing the Poisson Clicks Task.

**Figure 7.**
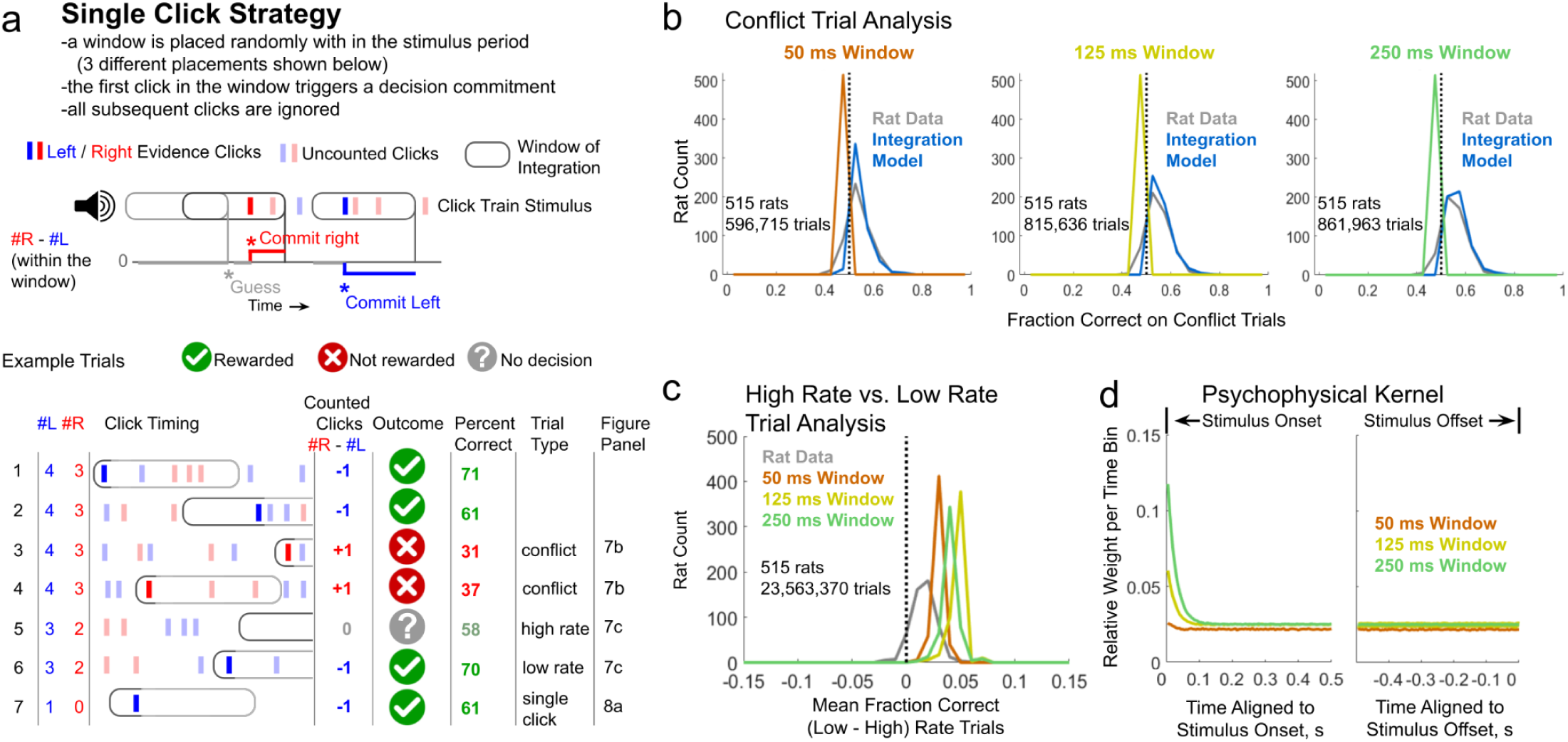
Analysis of performance of the Single Click Strategy on the Poisson Clicks task. **a)** Upper: Schematic describing the single click strategy. A window of integration begins at a random time within the stimulus period (here 3 possible placements are shown which lead to different outcomes). The first click within the window triggers a decision commitment. All other clicks are ignored. Lower: Example trials demonstrating how the single click strategy performs given different stimuli. Trials where the model samples no clicks result in a 50/50 guess. The outcome indicates how the single click strategy performs given the particular window placement shown while the percent correct indicates the mean performance of the single click strategy across all possible window placements on these example trials. The trial types refer to later panels in this and subsequent figures, b) Conflict trials are those the single click strategy is defined to get wrong. Here we test three variants of the single click model: 50, 125, and 250 ms windows of integration. Conflict trials are those with particular click times such that the window of integration is more likely to sample an incorrect click (see **(a)** example trials 3 and **4).** Histogram of th mean performance on the identified conflict trials plotted across rats for the single click strategy, integration model, and the rats’ data. Rats and the integration model perform better than chance on the conflict trials, consistent with them not using the single click strategy to perform the task. Only rats with at least 20 conflict trials are included in this analysis, **c)** Analysis of rat and model performance on trials with higher and lower overall click rates. First, trials are divided into groups with each particular # left and # right clicks. Second, each of those groups is split based on if the stimulus duration is greater or less than the group’s median. Shorter duration trials are flagged as “high rate trials” and longer duration trials are flagged as “low rate trials” (see **(a)** example trials 5 and **6).** Histogram plotted across rats of the difference in fraction correct on low versus high rate trials. In general rats perform slightly better on low rate trials. The single click strategy predicts a larger difference in performance since low rate (long) trials have less silence before the stimulus starts and therefore there’s less chance the window will sample silence leading the model to guess, d) Psychophysical kernels generated from 3 variants of the single click model run on trials at minimum 500 ms long generated with the same statistics as the trials rats performed. Trials are aligned to either the stimulus onset, left plot, or offset, right plot. For windows of integration longer than the mean inter click interval, the single click model over samples the first click, after which the strategy uniformly samples all clicks regardless of the window of integration duration.

As a first test we identify conflict trials the single click strategy is predicted to get wrong (Figure 7b). Just as we showed with the snapshot strategy, here too in the single click strategy, clicks that cluster have a lower probability of being sampled which can lead to an incorrect response being more likely. However because the single click strategy does not integrate at all there is a second means by which click timing can lead to wrong decisions. Since the model commits to the first click within the window of integration, a click placed close before a second click can mask it. The probability of a click being sampled is both a function of the duration of the window of integration and the inter-click interval that precedes the click. In the example conflict trials shown in Figure 7a (example trials 3 and 4) the inter-click intervals preceding the right clicks are on average larger than the intervals preceding the left clicks. Therefore there are more places the window of integration can begin that will sample a right click versus a left click, despite there being fewer right clicks. In Figure 7b we plot the performance of three variants of the single click model (50, 125, and 250 ms windows of integration) on conflict trials compared to the integration model and the rats’ performance. Rats in general (50 ms window of integration: 80.2%, 125 ms window of integration: 85.8%, 250 ms window of integration: 88.4%) perform significantly better than chance on conflict trials, consistent with them not using a single click strategy to perform the Poisson Clicks Task.

Similar to what we found with the snapshot strategy, the single click strategy also predicts improved performance on low rate compared to high rate trials (Figure 7c) for a similar reason. As done previously, we divide the trials into two groups, where each group contains an equal representation of trials with each particular number of right and left clicks, but differ in the rate at which those clicks are presented (see burst strategy section above for more details) As clicks spread out, the window of integration has a greater chance of sampling a click, and on average is then more likely to get the trial correct. While the rats do show a small improvement, it is smaller than the magnitude predicted by the single click model (Figure 7c).

Unlike the snapshot and burst strategy, which could only uniformly sample clicks throughout the stimulus period under very particular circumstances, the single click strategy, aside from the potential to over sample the first click, is guaranteed on average to uniformly sample all subsequent clicks. Following this early period, regardless of the duration of the window of integration, clicks have a uniform probability of being sampled regardless of when they occur during the stimulus period. The oversampling of the first click happens because there is no preceding click to mask it. The oversampling only occurs when the window of integration is longer than the average inter-click interval. For the majority of the rats in this data set, the first click is an uninformative stereo click, and oversampling it has no overall consequence in biasing the final decision, but it would still be detectable as a spike in the psychophysical kernel.

The single click strategy is theoretically capable of performing well on single click trials without any integration on multi-click trials (unlike the snapshot strategy) while maintaining a flat psychophysical kernel (unlike the burst strategy). To match the average rat performance on single click trials would require a ∼422 ms window of integration utilizing the single click strategy. However, most trials with a single click still contain a stereo click at the beginning of the stimulus. This limits the ability of the strategy to sample the non-stereo click. As done with the other degenerate strategies tested above, we fit an extended version of the single click model to the rats’ choice data (see Table 1 for the parameters that are included in this model). In Figure 8a we demonstrate that the variants of the single click model discussed in Figure 7 as well as the best fit single click model are not able to match the rats’ performance on single click trials. On average the best fit single click model is only able to perform at 53.4% correct on single click trials, compared with the rats’ mean performance of 65.6% correct. Of the 515 rats in this dataset 52 performed the frequency version of the Poisson Clicks Task which does not have an equivalent of an uninformative stereo click at the start of the stimulus, and 38 of those rats were presented at least 20 trials with only a single click and are included in this analysis. On average the single click model is able to perform better on these 38 rats’ single click trials, averaging 67.4% correct, compared to the rats’ 71.2% correct (r = 0.671, p < 0.01).

**Figure 8.**
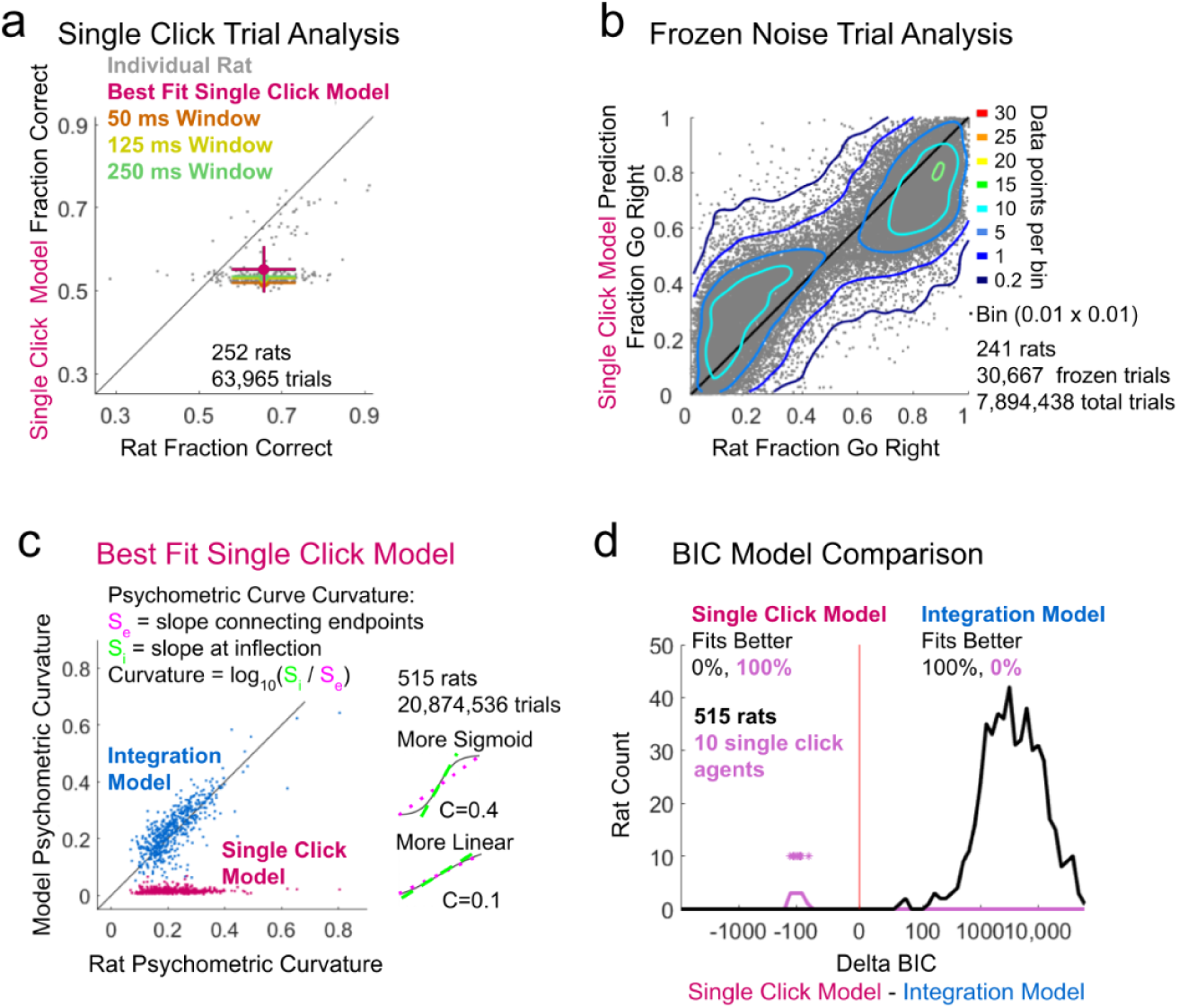
Analysis of the best fit Single Click Model to performance on the Poisson Clicks task. **a)** Rat and single click model performance on trials with only a single click (ignoring the stereo click that begins some trials). Individual dots represent individual rats that experienced at least 20 single click trials versus the best fit single click model. Mean performance of three variants of the single click model shown in color, b) Frozen noise trials are repeated trials with identical click train stimuli. The single click model is fit to all non-frozen trials, i.e. those that have unique stimuli, and tested on frozen trials. Data points represent an individual rat versus model predicted performance across a single unique frozen noise trial set. Scatter plot density is indicated with contour lines. Bins are 0.01 x 0.01 fraction go right, c) Analysis comparing the single click model and integration model psychometric curvature to the rats’ data. For each rat trials are separated into groups with an equal number of total clicks, for example, one group will contain all trials where the #R + #L = 10 clicks. Psychometric curves are then fit to each group’s data and curvature of the fit is calculated for each group. Curvature is defined as the log ratio of the slope connecting the psychometric curve’s endpoints (the value at the greatest positive and negative click difference) with the slope at the inflection in the middle of the curve (see inset). A linear psychometric will have a curvature of 0. Steeper sigmoid curves will have a greater curvature value. The mean curvature is calculated across all groups of trials for each rat. The best fit single click model (red points) results in psychometrics that are nearly linear, while the best fit integration model (blue points) results in curves that closely match the rats’ data. Each point represents data from an individual rat. d) Histogram of delta BIC analysis comparing the full integration model (10 parameters) with the full single click model (6 parameters). See Table 1 for a list of parameters included in each. No rats are better fit by the single click model (negative delta BIC). Synthetic choice data was generated using a range of parameters run on the single click strategy. All 10 of these synthetic agents (10k trials each) were better fit by the single click model. Individual synthetic data points also indicated with * as their histogram can be obscured.

Next we fit the single click model to all unique (non-frozen noise) trials, then test the prediction of the best fit model against the rats’ performance on frozen noise trials. In Figure 8b we plot the rats’ fraction choose “right” on each frozen noise trial set against the best fit single click model’s prediction. This degenerate strategy fails to accurately predict the rat’s choice behavior (compare Figure 8b to the integration model Figure 9e). On average the correlation between the rat’s performance and the single click model’s predicted responses is r = 0.881, the median difference between the predicted and actual performance being +/- 10.4%, and 9.7% of trials are poorly predicted having a greater than 25% difference between the predicted and actual response. For comparison the integration model has a correlation r = 0.951, median difference between predicted and actual performance of 4.4%, and 3.0% of trials are poorly predicted.

**Figure 9.**
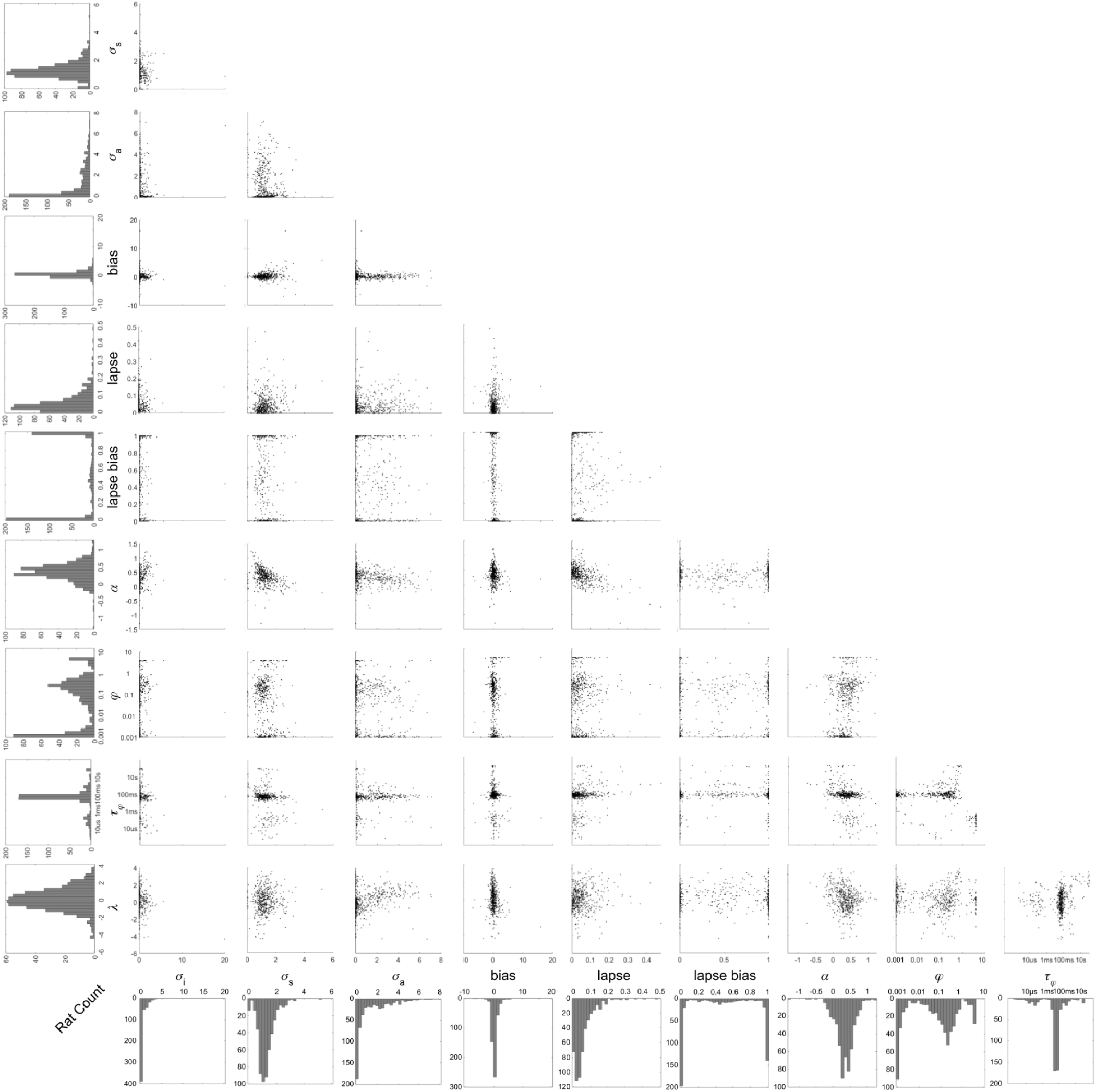
The best fit parameter values for the full integration model fit to 515 rats performing the Poisson Clicks Task. Each dot is a single rat. See Table 1 for a description of what role each parameter serves in the model.

Since the single click model only ever considers a single click, it is unable to replicate the sigmoid shape of the psychometric curve specifically when it’s plotted for trials with the same total number of clicks. Consider the set of trials with N total clicks; the probability of sampling any particular click on average is 1/N. Therefore the fraction of right responses would follow:

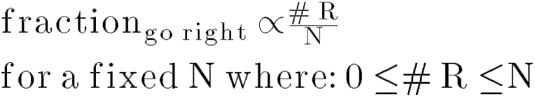

Where #R is the number of right favoring clicks. This is simply a line. While averaging across all possible trials can produce a sigmoid psychometric curve, when conditioned on a fixed total click count N the single click strategy is constrained to produce only linear psychometrics. To confirm this and test the rats’ data, we calculated the mean curvature of all psychometric plots for each rat separated by total click count.

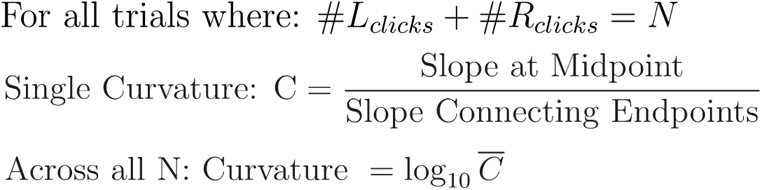

In Figure 8c we plot the mean curvature of the rats’ psychometric curves versus the curvature of the psychometric curves produced by the best fit integration model and best fit single click model. While the integration model is able to capture the curvature well (r = 0.590, p < 0.01), the single click model is not (r = 0) only producing linear psychometric plots with curvature ∼ 0.

Finally, we perform BIC analysis directly comparing the best fit single click model with the integration model, and find that 100% of rats (all 515) are better fit with the integration model. As done previously we generated 10 synthetic single click agents using a range of different parameter values and simulated their choices on the same 10,000 trials used earlier. When comparing the integration and single click model fits to these synthetic datasets we find the BIC analysis favors the single click model over integration for all of them (Figure 8d, purple), validating the model fitting and this analysis.

Once again we find the rats behavior is not consistent with them using the single click strategy to solve the Poisson Clicks Task. On average rats perform above chance on conflict trials (Figure 7b), trials with particular click timing such that the single click strategy is more likely to sample an incorrect click. Rats do not show an improvement in performance on low rate versus high rate trials at the magnitude expected if they were using a single click strategy (Figure 7c). Rats are able to perform significantly above chance on trials containing only a single click (Figure 8a) whereas the single click strategy performs close to chance (when there is a stereo click at the start of the stimulus period). When fit to unique trials the single click model is not able to accurately predict the rats’ response behavior on frozen noise trials (Figure 8b). The single click model is not able to generate sigmoid psychometric curves, like the rats do, when conditioned on trials with the same number of total clicks (Figure 8c). Finally the BIC analysis favors the integration model over the single click model for all 515 rats (Figure 8d).

### The Full Integration Model

Throughout this work we have demonstrated that the degenerate strategies tested are not able to fully account for all aspects of rats’ performance on the Poisson Clicks Task, and compared performance of the degenerate strategies to a “full integration” model (Figures 3b, 4d, 5b,d, 6d, 7b, 8c,d). The integration model used here is based on previously published work (Brunton et al., 2013; Scott et al., 2015) with a few minor changes. First, following the finding in (Brunton et al., 2013), we’ve eliminated the decision commitment bound parameter. This was a threshold if the accumulated value hit, a decision would be committed to and all subsequent clicks ignored. Previous analysis has shown the addition of this parameter does not significantly improve the fits and should be removed based on BIC analysis. By eliminating this parameter we are able to compute the model output for a given trial analytically without having to simulate the accumulated value distribution for each timestep, making the analysis of this large dataset more tractable.

The second change we made to the original integration model is the addition of two new parameters: lapse bias and *α* (Table 1, Figure 9). In the original model lapse was defined as the fraction of trials the model makes a random response, 50% left and 50% right, irrespective of the stimuli. Here the “lapse bias” parameter controls what fraction of the lapse trials have a left or right response.

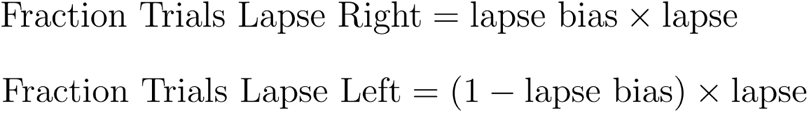

This allows the asymptotic tails of the psychometric curve to be asymmetric. BIC analysis supports the addition of this parameter in 344 of the 515 rats (66.8%). The 171 rats where BIC does not support the addition of this parameter, 133 (77.8%) are best fit with a lapse parameter value less than 5%, and therefore the lapse bias parameter has a negligible impact since there are few lapse trials to act on. Overall the best fit value for the lapse bias parameter follows a trimodal distribution (Figure 9): 30.5% of rats are best fit with a lapse bias ≥ 0.95 (meaning they almost always respond right when lapsing), 41.9% of rats are best fit with a lapse bias ≤ 0.05 (meaning they almost always respond left when lapsing), and the rest (27.6%) follow relatively normal distribution with mean 0.501 +/- 0.227 (standard deviation).

The second new parameter, *α*, defines how “scalar” the integrated values are relative to the noise (Fechner, 1860; Gibbon, 1977; Meck and Church, 1983). In the original model (Brunton et al., 2013) noise grows with the square root of the number of clicks while the integrated value grows linearly. In this regime the signal to noise improves with more clicks (i.e. 1 vs. 2 clicks, 3 total, will have a lower signal to noise than 2 vs. 4 clicks, 6 total) producing response behavior that is not scalar. Scalar means that the signal to noise is constant (Gibbon, 1977; Meck and Church, 1983) given a fixed ratio of right and left clicks, and therefore performance would be the same on all trials with the same ratio of right and left clicks regardless of the click total. Later work found rat response behavior in a visual version of the Poisson Clicks Task, Flicker (Scott et al., 2015), was better fit by a model that was scalar, where noise and accumulated evidence grew at the same rate. When constrained to these two options we find that 437 of the 515 rats (84.8%) are better fit by a nonscalar integration model (consistent with (Brunton et al., 2013)). However, rather than forcing the model to adopt one of these two options, we introduce the free parameter *α* which allows the model to be either, scalar as in (Scott et al., 2015), not scalar as in (Brunton et al., 2013), or anywhere in between. Here noise in the model grows with the square root of the number of clicks, but now accumulated evidence is given by:

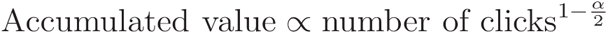

When *α* = 0 the accumulated value grows linearly with the number of clicks, as in (Brunton et al., 2013) resulting in non-scalar response behavior, and when *α* = 1 the accumulated values grows with the square root of the number of clicks (Nieder and Miller, 2003), the same rate as the noise, producing scalar behavior as in (Scott et al., 2015). Now with the free parameter *α*, the accumulator is not constrained to only these two regimes but can adopt a signal to noise behavior along a continuum. BIC analysis supports the addition of this parameter in 376 of the 515 rats compared to fixing *α* at 0, and 509 of the 515 rats compared to fixing *α* at 1. In Figure 9 we plot the best fit value of each parameter in the integration model against each other. On average rats are best fit with *α* = 0.344 with 95% of the rats having a value between -0.164 and 0.800. Of the 515 rats, 52 rats are best fit with *α* < 0. In this regime the accumulated number of clicks grows supralinearly, and therefore the signal to noise improves faster than in (Brunton et al., 2013). In contrast 6 rats are best fit with *α* > 1. In this regime the accumulated number of clicks grows slower than the noise; performance decreases given more clicks for a fixed click ratio, i.e. 2 vs. 4 clicks will have a lower signal to noise than 1 vs. 2 clicks.

### Testing the Full Integration Strategy

BIC analysis has demonstrated that the integration model provides a better fit to the rats’ choice data than any of the three degenerate strategies discussed above (Figures 4d, 6d, and 8d), however this does not indicate how good the overall quality of the fit the integration model provides is. Here we will apply the same analysis we used to test the degenerate strategies to the full integration model to better assess its ability to explain the rats’ choices on the Poisson Clicks Task.

Conflict trials are defined as those with particular click timing such that a strategy is more likely to sample incorrect clicks and get the trial wrong. The closer a strategy is to perfect integration the fewer conflict trials exist. Unlike the degenerate strategies that only sample a small portion of the stimulus on any given trial, and the results of the subsampling were the main driver of what defined the conflict trials, the full integration model always samples the entire stimulus. Despite this there are still means by which full integration can get particular trials wrong. First, the model can have a small bias favoring one response (bias and lapse bias parameters), leading to errors on trials with a small click count difference in the opposite direction. Second, click adaptation (*φ* and *τ*_*φ*_ parameters) can lead to the suppression of correct clicks, for example if they occur in a burst, resulting in an incorrect response. Finally, the *λ* parameter, the inverse of the integrator’s memory time constant, can cause the model to favor earlier or later periods within the stimulus (Usher and McClelland, 2001). The model will tend to get trials wrong that have more correct clicks in the portion of the stimulus it weighs less. In Figure 10a we plot a histogram of the rats’ mean performance on conflict trials identified from the best fit integration model. A majority of the rats (57.9%) do perform below chance on these trials, consistent with the integration model’s performance. Rats that continue to perform above chance on conflict trials are an indication that in these instances we are either not modeling bias, click adaptation, and the integrator’s time constant correctly, or more likely we are missing additional mechanisms by which an integrator can make mistakes.

**Figure 10.**
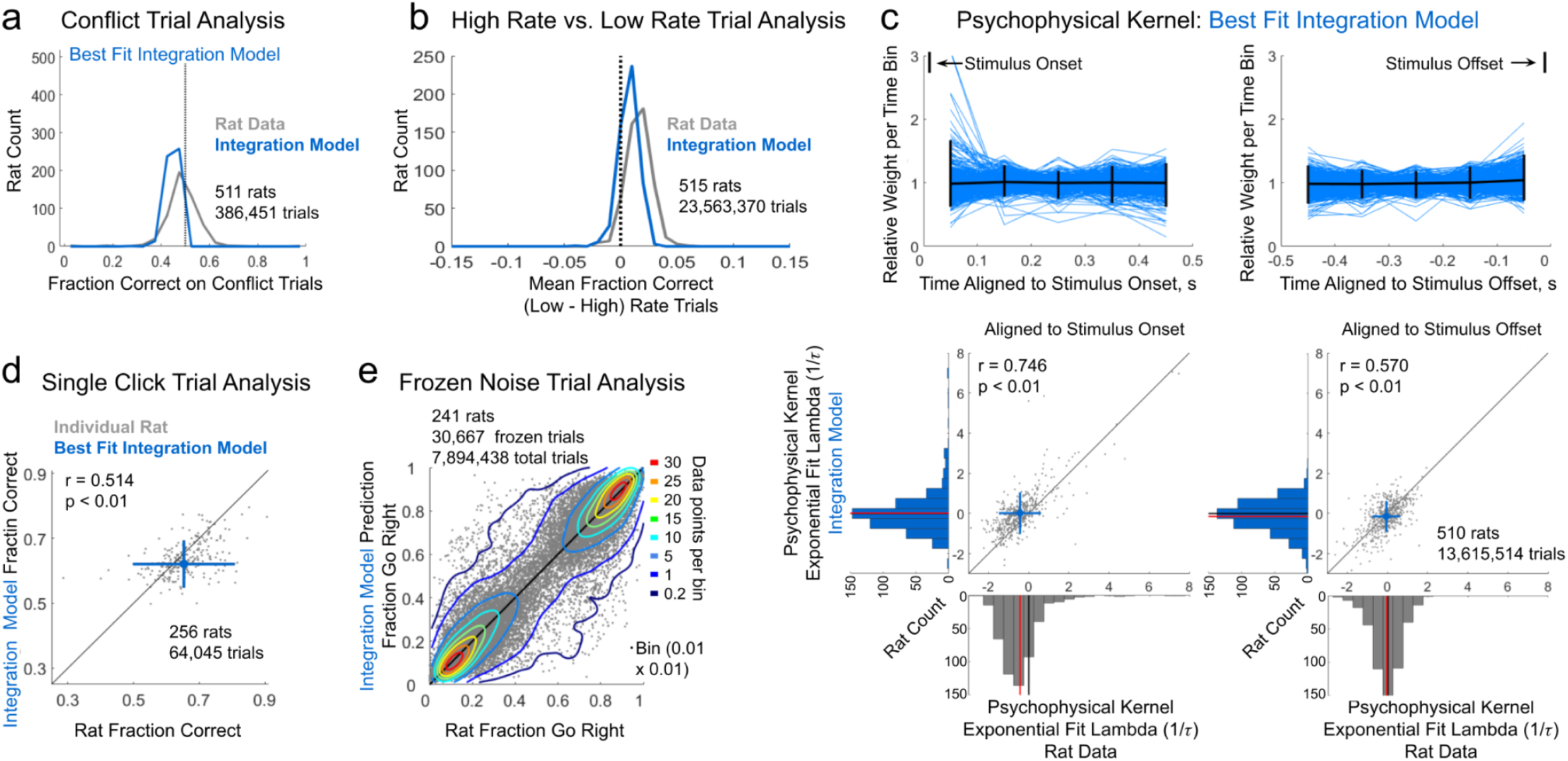
Analysis of the best fit Integration Model to performance on the Poisson Clicks task. **a)** Conflict trials identified as those the best fit integration model is predicted to get wrong. Conflict trials are those with particular click times such that the integration model under counts the clicks favoring the correct response. Histogram of rats’ fraction correct (gray) and the integration model’s fraction correct (blue) on conflict trials. Only rats with at least 20 conflict trials are included in this analysis. 57.9% of rats perform on average below chance on conflict trials, **b)** Analysis of rat and model performance on trials with higher and lower overall click rates. First, trials are divided into groups with each particular # left and # right clicks. Second, each of those groups is split based on if the stimulus duration is greater or less than the group’s median. Shorter duration trials are flagged as “high rate trials” and longer duration trials are flagged as “low rate trials”. Histogram plotted across rats of the difference in fraction correct on low versus high rate trials. In general rats perform slightly better on low rate trials similar to the integration model fits, **c)** Top: The psychophysical kernel produced by the best fit integration model to each individual rat (blue) aligned to stimulus onset (left) or offset (right) for trials with stimuli longer than 500 ms. Mean shown in black. Vertical bars indicate 95% of the rats. Bottom: An exponential function is fit to each psychophysical kernel produced by the best fit integration model for each rat. Plot of the lambda value (1 A) from this exponential fit compared to the lambda of the fit to each rat’s psychophysical kernel (Figure 2c). Left aligned to stimulus onset, right aligned to stimulus offset. The integration model results in psychophysical kernels that are similar to the rats’ data. Mean plotted in blue. Horizontal and vertical bars indicate 95% of the rats. Histograms of the lambda values plotted separately for the integration model and rats’ data. Rat data histograms are the same as in Figure 2c. **d)** Rat and integration model performance on trials with only a single click (ignoring the stereo click that begins some trials). Gray dots represent individual rats that experienced at least 20 single click trials versus the best fit integration model. Mean performance plotted in blue. Horizontal and vertical bars indicate 95% of the rats, **e)** Frozen noise trials are repeated trials with identical click train stimuli. The integration model is fit to all non-frozen trials, i.e. those that have unique stimuli, and tested on frozen trials. Not all rats experienced frozen noise trials when performing the Poisson Clicks task. Only rats that were presented at least 40 different frozen noise trials, each at least 100 times, are included in this analysis. Data points represent an individual rat versus model predicted performance across a single unique frozen noise trial set. Scatter plot density is indicated with contour lines. Bins are 0.01×0.01 fraction go right.

Next, as done previously, we divide the trials into two groups, where each group contains an equal representation of trials with each particular number of right and left clicks, but differ in the rate at which those clicks are presented (see burst strategy section above for more details). As clicks spread out, click adaptation has a smaller effect on the final integrated value leading to more accurate performance. Rats do show a small improvement in performance on low rate compared to high rate trials, consistent with the magnitude of the effect expected from the integration model (Figure 10b), and previous findings (Brunton et al., 2013).

One hallmark of the rats’ performance on the Poisson Clicks Task is their relatively flat psychophysical kernels, indicating they weigh or sample evidence across the entire stimulus uniformly (Figure 2c). The integration model is not only able to capture this overall uniform sampling behavior well (Figure 10c top), but also fit the individual rats’ particular kernel profiles (Figure 10c bottom).

Trials that contain only a single (non-stereo) click were a challenge for the degenerate strategies to perform well on. The integration model by contrast (Figure 10d) is able to more accurately match the rats’ performance (r = 0.514, p < 0.01). Here we include data from all rats that were presented at least 20 single click trials.

Finally, we use the frozen noise trials to examine how well the integration model is able to predict the rats’ response behavior (Figure 10e). Here we fit the integration model to all unique (non-frozen trials) then plot the model’s predicted response on each frozen trial set against the rats’ actual fraction respond “right”. On average the correlation between the rat’s performance and the model’s predicted responses is r = 0.951 with the median difference between the predicted and actual performance being +/- 4.4%. Overall the integration model offers a better prediction of the rats’ choice behavior than any of the degenerate strategies explored earlier (compare to Figures 4d, 6d, and 8d). A small fraction of the frozen noise trials are poorly fit by the model as indicated by the data points that lie far from the diagonal (3% of trials have a difference between prediction and data greater than 25%). This indicates there must be some large component of the rats’ decision making process that is not captured on a small but statistically significant fraction of the trials.

Unlike the degenerate strategies examined above, the full integration model is able to fit many aspects of the rats’ response behavior well: the majority of rats do perform below chance on conflict trials; rats do show a small increase in performance on low rate trials compared to high rate trials, consistent with the integration model; the integration model is able to fit the flat psychophysical kernels extracted from the rats’ response data; full integration is able to match the rats’ performance on single click trials; and the integration model provides the most accurate prediction of the rats’ choices on frozen noise trials.

As a final test we wanted to analyze a set of trials that only a full integration strategy could perform well on. We refer to these trials as “bookend trials”. These are trials where the number of right and number of left favoring clicks differs by only 1, and where the first and last click favor the same response (ignoring any initial stereo click). On correct bookend trials the first and last click favor the correct response (Figure 11a). Here any strategy that ignores early or late clicks will end up sampling an equal number of left and right clicks, leading to guesses. Only a strategy that integrates the early and late periods of the stimulus, either by integrating the entire stimulus or simply ignoring the middle portion of the stimulus, can consistently perform above chance on these trials. At the bottom of Figure 11a we plot a histogram of the mean performance across rats on the correct bookend trials they experienced. The majority of rats perform above chance on these trials, consistent with the full integration model. A strategy that ignores the middle portion of the stimulus also performs above chance on these trials.

**Figure 11.**
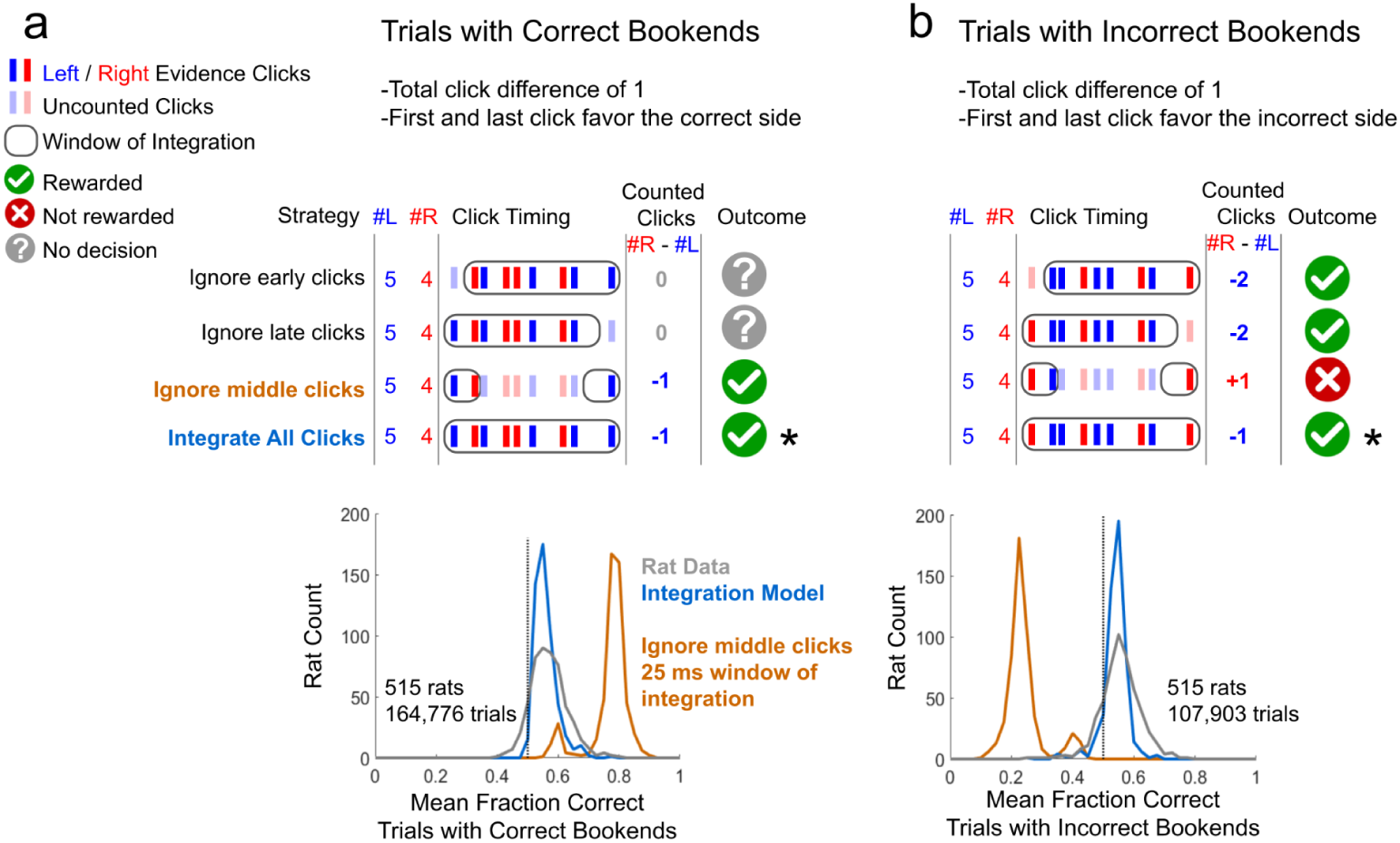
Analysis of rat and model performance on trials with bookend clicks. **a)** Correct bookend trials, defined as those with a total click difference of 1 and where the first and last click favor the correct response (ignoring the stereo click that begins some trials). 4 possible strategies are shown. Integrating the entire stimulus or only integrating the early and late periods of the stimulus leads to a correct response on these trials. Histogram of each rat’s average fraction correct on correct bookend trials along with the best fit integration model’s performance on these trials. The majority of rats and the best fit integration model perform above chance on correct bookend trials. Mean performance of an agent that ignores middle clicks and only considers clicks within the first and last 25 ms of the simulus is included as well, **b)** Incorrect bookend trials are like correct bookend trials except the first and last click favor the incorrect response, again ignoring any stereo clicks. The same 4 strategies are demonstrated along with a histogram of each rat’s average fraction correct on incorrect bookend trials along with the best fit integration model’s performance. The majority of rats and the best fit integration model perform above chance on incorrect bookend trials. Mean performance of the same “ignore middle clicks” agent from (a) is included here. Only integration of the entire stimulus leads to performance that is better than chance on both trial types (marked with an *).

While we can easily rule out the “ignore middle clicks” strategy since it would result in a psychophysical kernel with large spikes at the earliest and latest time bins, we can also look at a complementary set of trials, the incorrect bookend trials (Figure 11b). While a full integration strategy continues to perform above chance on these trials, now ignoring the middle portion of the stimulus leads to performance significantly below chance. The majority of rats on average perform above chance on incorrect bookend trials, again consistent with performance of the full integration model. Only a full integration strategy is able to perform above chance on both correct and incorrect bookend trials.

## Discussion

Understanding the strategy subjects use to perform cognitively demanding tasks is critical to properly interpreting the data they generate. If a task is designed to explore a particular cognitive process, but subjects are able to perform well on that task without engaging that process, additional tests and analysis must be run to confirm which strategy the subjects are using. The ability to gradually accumulate noisy perceptual evidence, and use it to form a decision, is a cognitive process that has been the focus of extensive investigation for many years (Brody and Hanks, 2016; Carandini and Churchland, 2013; Glickman and Usher, 2019; Gold and Shadlen, 2000; Hanks and Summerfield, 2017; Huk and Shadlen, 2005; Keung et al., 2019; Newsome and Paré, 1988; Odoemene et al., 2018; Shadlen and Newsome, 1996; Znamenskiy and Zador, 2013). While the optimal strategy to perform these tasks requires one to integrate information from the entire stimulus presented, other degenerate strategies exist that allow a subject to perform well without integrating (Ditterich, 2006; Hyafil et al., 2023; Pinto et al., 2018; Stine et al., 2020). Here we compile the largest dataset of animals performing a perceptual decision-making task (Kopec et al., 2024), the Poisson Clicks Task (Brunton et al., 2013; Sanders and Kepecs, 2012), that requires the accumulation of noisy perceptual evidence, and perform a series of analyses to determine if the rats are using any of these degenerate strategies, or fully integrating the stimulus. The discrete pulsatile random nature of the stimuli in the Poisson Clicks Task is integral in being able to determine what strategy subjects use to perform it. Here we will briefly review these findings highlighting how features of the Poisson Clicks Task stimuli facilitate strategy identification.

The Poisson Clicks Task can have trials with as few as a single click. In general, the degenerate strategies are unable to perform well on such trials while maintaining a flat psychophysical kernel. If the burst strategy is able to commit to a decision based on a single click, then on trials with many clicks, the early clicks will trigger decision commitment and the later clicks will be ignored. This would result in a steeply decaying psychophysical kernel (Figure 3d). For the snapshot strategy to perform well on single click trials, a snapshot on the order of 100s of milliseconds is required. On trials with many clicks, such a long window of integration would result in boundary effects, as the earliest and latest periods of the stimulus would be sampled less than the middle portion (Figure 5d). The single click model can perform well on trials with only a single click as long as there is no stereo click at the onset of the stimulus. However, on trials with no stereo click, windows of integration longer than the mean inter-click interval result in an oversampling of the first click and a spike in the psychophysical kernel (Figure 7d).

While the click train stimuli are generated with a specific underlying total click rate, due to the Poisson nature of the click timing, trials will have an actual click rate higher or lower than the true generative rate. Trials with higher than average click rates are more likely to contain a burst of clicks. Therefore, the burst strategy predicts better performance on these trials as trials with no burst are answered randomly with a guess. The snapshot and single click strategies predict the opposite. Trials with higher click rates have the same number of clicks as trials with lower click rates but these clicks are compressed into a shorter duration stimulus. This results in a longer period of silence prior to the stimulus onset, a longer period where the snapshot and single click strategies would sample no clicks if the window of integration happened to be placed there. This longer period of silence increases the likelihood the snapshot and single click strategies would guess, decreasing their overall performance relative to lower rate trials.

The highly varied timing of clicks within the stimulus occasionally result in trials that trick different strategies into getting the trial wrong. We refer to these as “conflict” trials because the choice the strategy favors conflicts with the presented evidence (previously referred to as “disagree trials” see (Hyafil et al., 2023; Levi et al., 2018)). These are not a universal set of trials but specific to the strategy being tested. What is a conflict trial for one strategy may not be for another. For example, if a trial contains more left clicks but the fewer right clicks are clustered in a burst, the burst strategy would respond to the burst of right clicks and get that trial wrong. On the other hand, if the more left clicks were clustered together the snapshot and single click strategies would be less likely to sample them, leading to an increased probability of an incorrect response. We know the rats performing the Poisson Clicks Task are not perfect integrators, and different strategies predict different patterns of mistakes. Therefore, if we identify a strategy that predicts mistakes on the same trials the subjects get wrong, that’s strong evidence the subjects are using that strategy. However, the majority of rats perform above chance on the conflict trials identified for all of the variants of the three degenerate strategies tested here, suggesting that they are not using any of those strategies to perform the Poisson Clicks Task.

Finally, because of the nature of the Poisson Clicks Task, stimuli predict quantitatively different performance between degenerate and integration strategies as outlined above. Given the large number of trials performed, simple model fitting and BIC analysis is sufficient to determine which is more likely to be closely aligned with the strategy the subjects are using.

This is in contrast to other integration of evidence decision-making tasks where evidence is presented continuously, such as the Random Dot Motion Task (Gold and Shadlen, 2000; Newsome and Paré, 1988; Shadlen and Newsome, 1996), a visual accumulation of evidence task or the Cloud of Tones Task (Znamenskiy and Zador, 2013), an auditory accumulation of evidence task. First, in continuous evidence tasks there are no periods when no information is being presented; because of this, snapshots of arbitrarily short duration will still capture information and can uniformly sample the stimulus period. Second, while the strength of the information presented can be adjusted, for example, by changing the coherence with which the dots move or the fraction of tones in the correct band, the overall rate at which the information is presented is generally constant. There is no equivalent of high and low rate trials that highlight how the degenerate strategies behave differently under these regimes. Third, continuous evidence integration tasks typically do not have trials where only a single bit of information is presented. While these trials are also rare in this dataset, constituting ∼0.2% of all trials, they provide a powerful constraint on which strategies could successfully explain all aspects of the rats’ response behavior. Since continuous evidence accumulation tasks lack these features: periods of silence between bits of evidence, high and low rate trials, and trials with only a single bit of evidence, the analyses performed here, which allows one to rule out specific degenerate strategies without altering the task structure, could not be applied.

Continuous evidence integration tasks also tend to present multiple bits of information simultaneously. In the Random Dot Motion Task many dots are presented on each frame of the video, some number of which are moving coherently, and in the Cloud of Tones Task, different tones play at random constantly overlapping each other. This adds an extra layer of complexity to modeling how a subject deals with this large amount of simultaneous information. For example, are they averaging it all together, which is its own form of integration? Are subjects subsampling a small portion of the information, essentially taking a snapshot in stimulus space? Are they utilizing only the most extreme portions of the stimulus at each point of time? Or, are subjects randomly selecting only a single bit of the many bits presented to use at each timepoint? In contrast, discrete evidence tasks only present one bit of information at a time, greatly simplifying the modeling of decision making.

While discrete evidence integration tasks have a number of advantages, there are changes that can be made to the Poisson Clicks Task to improve it. First, while a fixed generative Poisson rate will lead to stimuli with different actual rates, making trials using different generative rates, e.g. 10 Hz, 20 Hz, and 40 Hz presented to the same subject, will sample the stimulus space better. Second, trials with only a single click serve as a key piece of evidence when fitting models to a subject’s data, and therefore the frequency of their occurrence should be increased. Single clicks tend to occur on the shortest duration trials and therefore the clicks do not uniformly sample the stimulus period but rather tend to occur near its end. Trials could be engineered to avoid this. Finally, in the frequency version of the Poisson Clicks Task we explicitly ensure high and low pitch clicks do not overlap. However in the location version of the task each click train is independent and clicks can occasionally overlap (∼10.8% of all clicks overlap with another click from the same or opposite speaker on trials with a total rate of 40 Hz). When this happens, the sound pressure waveforms can constructively and destructively interfere in complex ways. Eliminating these “collisions” would remove an unknown effect from the decision-making process. Some rats in this dataset (9.9%) performed the Poisson Clicks Task with stimuli that are selected to avoid any overlap of clicks (see Methods: Dataset).

Previous work (Stine et al., 2020) has shown that the more trials a subject performs, the easier it becomes to determine which strategy they are using to solve an accumulation of evidence task. Their analysis determined that ∼12,000 trials are sufficient to reliably distinguish between strategies. In the dataset we present here, the minimum number of non-violation trials a rat must perform to be included is 10,000 with the actual mean per rat being ∼70,000. While this many trials may be difficult if not impossible to obtain from individual human subjects, rodents in a semi-automated training environment can easily generate this much data in a few weeks.

The trials included in this data set were all generated at random following Poisson statistics. While this maximizes sampling the stimulus space and is sufficient to generate enough trials to distinguish the strategies tested here, there’s no reason engineered trials could not be designed to specifically test different strategies. Engineered conflict trials could be designed to maximize the performance difference, allowing one to differentiate which strategy a subject is most likely using, with far fewer trials.

Despite providing a better fit to the rats’ choice data compared to the degenerate strategies examined here, the integration model is clearly not sufficient to account for all possible idiosyncrasies in an individual rat’s performance. The model’s predicted performance on frozen noise trials, while highly correlated with the rats overall performance, demonstrates there is ample room for improvement. Our approach of repeatedly presenting trials with frozen noise assures that deviations in the animal’s performance on these trials from the model prediction can be determined with a high degree of accuracy. The trials the model is least able to accurately predict are those where the rat responds close to chance, i.e. these are cognitively the most difficult, and therefore potentially the most interesting trials to study.

Additionally, the integration model as formulated here cannot account for trial history effects, which are a small but significant factor in determining a subject’s response on perceptual decision-making tasks (Busse et al., 2011; Gold et al., 2008; Hermoso-Mendizabal et al., 2020; Morcos and Harvey, 2016; Scott et al., 2015) including the Poisson Clicks Task (Gupta et al., 2024). Second, recent work has demonstrated that accounting for shifts in a subject’s internal state (Ashwood et al., 2022; Calhoun et al., 2019), between periods when they’re highly engaged in the task contrasting with periods when their responses are less driven by the stimulus, can significantly improve the accuracy with which a model can predict the subject’s responses. Finally we must concede there are factors we have not even thought to consider which may significantly improve a model’s predictive power. The large number of subjects included in this dataset should allow for the identification of factors that affect only a small fraction of the individuals, while the large number of trials should allow for the identification of factors that have small but significant effects overall. For these reasons we hope others will find this dataset compelling enough to test hypotheses against.

As scientists we must always be willing to question the conclusions we draw from our data (Palminteri et al., 2017; Wilson and Collins, 2019). Cognitive neuroscience is built on a foundation constructed from a panoply of different behavioral tasks (Thorndike, 1911). The insights we glean from these experiments is predicated on knowing how the subjects are performing them (Brunton et al., 2013; Ditterich, 2006; Hyafil et al., 2023; Pinto et al., 2018; Stine et al., 2020; Waskom and Kiani, 2018). Here we have compiled one of the largest perceptual decision-making data sets, in an effort to critically evaluate the subjects’ performance, and understand the strategy they use to perform the task. While we have demonstrated, using multiple independent analysis, that a full integration strategy better explains the rats’ choices compared to the three degenerate strategies tested here, we remain open to the idea that an as yet unknown strategy may still fit better (Wilson and Collins, 2019). Therefore, to help further our understanding of how the cognitive process of accumulating noisy perceptual evidence to form a decision functions, we hope others will find this data set to be a useful and fruitful test bed.

## Acknowledgements

We would like to thank all members of the Brody Lab, past and present, who helped make this dataset possible, most especially our animal handling technicians who worked with the rats on a daily basis. We thank Alexandre Hyafil for fruitful discussions surrounding this work. We would like to thank the IT support team at the Princeton Neuroscience Institute, especially John Wiggins, Randee Tengi, and Garrett McGrath, who helped ensure the computers and network infrastructure on which these experiments relied were always operational, as well as Gary Drozd, Todd Antonakos, and Kevin Durham for their help in maintaining the research space over the years. Finally, we are grateful to the animal husbandry and veterinary staff at Princeton University, including Laura Conour, Jamus MacGuire, Grace Barnett, Susie Chow, Brian Ludwig, Paul Cunningham, Kirsten Gerhart, and their entire team who helped care for and ensure the well-being of the animals in this study. This work was supported by the Howard Hughes Medical Institute and the National Institutes of Health.

## Methods

### Subjects

A total of 515 male Long-Evans rats (*Rattus norvegicus*) between the ages of 2 months and 24 months were used for this study. Rats did not begin water restriction and training until their weight exceeded 225 g. The majority of rats in this study were pair housed. Animal use procedures were approved by the Princeton University Institutional Animal Care and Use Committee (IACUC) and carried out in accordance with National Institutes of Health standards.

### Behavior

Rats were trained to perform an auditory evidence-accumulation decision task, the Poisson Clicks Task. For each session, rats were placed in a behavioral training box that itself was located within a sound attenuation chamber with active ventilation. The behavioral training box has three conical nose ports and two speakers (see Figure 1a for arrangement). Each nose port is equipped with an infrared beam to detect nose poke events and a white LED that can illuminate the port. The left and right ports are equipped with a sipper tube and a solenoid valve (Lee Valve Company, LHDA1231115H) that dispenses distilled water from a 10 gallon gravity fed tank. The center port has an enlarged opening in the center to facilitate the rat’s ability to hold their nose in the port.

Once a rat’s mass passed 225 g they were placed on a water restriction schedule and training began. Whenever possible training would occur at the same time everyday, in the same behavioral training box, seven days per week. Training sessions typically lasted between 1.5 and 3 hours. Animals that trained between 8pm and 8am were housed on a reverse light cycle. Supplemental water was offered in two ways: 1) rats were returned to their home cages after a training session, where food was available *ad libitum*, and offered 1 hour free access to water via the sipper tube in their cage; 2) rats were placed in a modified behavioral training chamber with one nose port where water consumption could be monitored. Rats in this second group were restricted to receive a specific percentage of their body mass in water (between 3 - 5%). Any amount not earned while training would be offered in the supplemental period post training. Rats were free to earn more than their restricted allotment by performing more trials while training. Rats in this second group had food available *ad libitum* in their home cage and while training but not when receiving their supplemental water.

The behavioral training box was either controlled via a National Instruments Data Acquisition (NI DAQ) card connected to a computer running a real-time Linux kernel, or a Bpod 2.3 (Sanworks). Both systems were managed by a Windows computer running Bcontrol, a custom matlab behavioral training software package (see https://brodylabwiki.princeton.edu/bcontrol/index.php?title=Main_Page for a description of how Bcontrol works and https://github.com/Brody-Lab/ExperPort_public to download the full code package). For NI DAQ controlled rigs sounds were generated via a Lynx sound card on the linux computer, while on Bpod rigs sounds were generated from an Analog Output Module (Sanworks). In both cases the signal was amplified by a mini class D stereo amplifier (Leapai LP-2020AD) and played through a pair of planar tweeters (Bohlender Graebener Neo3W).

### Training

Rats in this dataset, compiled over more than a dozen years, did not learn the Poisson Clicks Task using the exact same training pipeline. However, the training in general followed the training stages outlined below. Training on the frequency and location versions of the Poisson Clicks Task followed the same training protocol.

1. In this stage the rat learns to poke in the left port. Left port is illuminated. Rat receives a 24 µl drop of water if they poke into the left port. At the end of the session if the rat has completed at least 50 trials at 80% correct advance to the next stage.
2. In this stage the rat learns to poke in the right port. Right port is illuminated. Rat receives a 24 µl drop of water if they poke into the right port. At the end of the session if the rat has completed at least 50 trials at 80% correct advance to the next stage.
3. In this stage the rat learns to poke in the center port before poking in a side port. Center port is illuminated. Rat must poke into the center port after which one of the side ports illuminates. Rat must poke into the illuminated side port to earn a water reward. Trials are presented in blocks of 20 alternating sides. At the end of the session if the rat has completed at least 50 trials they advance to the next stage.
4. In this stage the rat learns to respond to auditory stimuli rather than simply poking in whichever port is illuminated. Same as stage 3 except now sounds play and the side ports no longer illuminate. The center poke initiates a train of clicks with gamma = +/- 4 that continues to play until the rat makes a correct side poke. Trials are presented in blocks of 20 alternating sides. At the end of the session if the rat has completed at least 150 trials at 70% correct advance to the next stage.
5. In this stage the rat must learn which auditory stimulus indicates a left vs a right trial. Same as stage 4 except trials are presented in blocks of 5 alternating sides, and after trial 40 the correct side is randomly selected. At the end of the session if the rat has completed at least 150 trials at 70% correct advance to the next stage.
6. In this stage the rat learns to hold their nose in the center port for the fixation period. The center poke duration begins at 1 ms the first day the rat starts this stage and grows gradually. Every trial the rat is able to complete without violating grows the fixation duration by 1 ms. A violation is defined as any break in fixation that exceeds 5 ms. A violation event replays the same trial until the rat is able to complete it. At the end of the session, if the fixation duration has grown to 1.5 s advance to the next stage, if not they resume where they left off on the following day.
7. In this stage the rat must perfect their ability to hold fixation and respond to the easiest stimuli. At this point the rats are doing the full Poisson Clicks Task with a 1.5 s fixation period, but only with the easiest trials, gamma = +/- 4, and all stimuli begin when fixation begins and play for 2 s. At the end of the session, if the rat has completed at least 120 trials at 75% correct advance to the next stage.
8. In this stage the rat is introduced to a range of different trial difficulties. At the start of this training stage the presented gammas are expanded to +/- [4, 5]. At the end of the session if the rat completed at least 120 trials at 70% correct the gammas are set to +/- [3, 5].
9. In this stage the rat learns to perform the task with shorter duration stimuli. Rats continue to perform the task with trial gammas = +/- [3, 5]. Now the duration of the stimulus will be reduced. At the start of this stage the stimulus duration is set to end at the end of fixation, 1.5 s duration stimuli. As the duration of the stimulus is reduced the period of silence at the beginning of fixation grows. At the end of each session if the rat has completed at least 120 trials at 70% correct the duration of the stimulus is reduced by 100 ms. Once the duration is reduced to 1 s advance to the next stage.
10. In this stage the rat is introduced to stimuli that have a range of different durations. From here on the longest stimuli are kept fixed at 1 s. At the end of each session if the rat has completed at least 120 trials at 70% correct the minimum stimulus duration is reduced by 100 ms. A trial’s stimulus duration is selected from a uniform distribution between the minimum duration and 1 s. Once the minimum duration has been reduced to 200 ms advance to the next stage.
11. In this stage the rat must perfect performance on trials with the full duration range. The stimuli gammas are set to +/- [2, 3, 4, 5]. At the end of the session if the rat has completed at least 150 trials at 70% correct advance to the next stage.
12. In this stage the stimuli gammas are gradually reduced making the set of trials presented more difficult. At the start of this training stage the easiest trials have a gamma = +/- 5 and the most difficult trials have a gamma = +/- 2. At the end of the session if the rat has completed at least 120 trials at 70% correct the gammas are reduced by 0.1. Once the gammas have reached +/- [1, 2, 3, 4] advance to the next stage.
13. In this stage a violation event no longer replays the same stimulus. From this stage on a violation is paired with a 1 s time out and a 1 s burst of noise from the speakers and an entirely new trial is initiated. In this way the rat cannot sample the same stimulus multiple times. At the end of each session if the rat has completed at least 150 trials at 70% correct the gammas are reduced by 0.1. Once the gammas reach +/- [0.5, 1.5, 2.5, 3.5] advance to the next stage.
14. This is the final stage where the rat is performing the full Poisson Clicks Task with the full range of stimulus gammas +/- [0.5, 1.5, 2.5, 3.5] and the full range of stimulus durations (200 ms to 1 s).

### Behavior Analysis

A psychometric curve is fit to the rat’s response behavior:

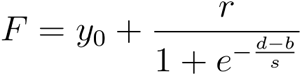

where F is the fraction respond “right”, y_0_ is the lower asymptote, y_0_ + r is the upper asymptote, d is the click difference (#R - #L), b is the position of the curve’s inflection point along the horizontal axis, and s along with r control the steepness of the curve at the inflection point (see Slope below).

Bias, the click difference that results in an equal probability of making either response is given by:

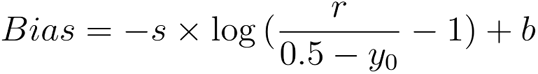

The slope at the psychometric curve’s inflection point, the change in fraction correct per click is given by:

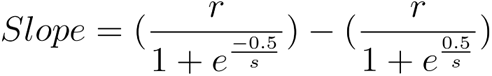

### Model Fitting

All models were run in Matlab 2019b on Windows computers. Each model was fit to each individual rat’s choice data starting from 5 independent random seeds. For the parameters that were not fixed in value, seeds were selected from a uniform distribution within the following range: *σ*_i_ [0, 5], *σ*_s_ [0, 4], *σ*_a_ [0, 20], bias [-10, 10], lapse [0, 0.4], lapse bias [0, 1], *α* [0, 1], *φ* [-1, 0.02], *τ*_*φ*_[0, 2], window of integration [10, 1000] (milliseconds), *λ*[-1, 1]. Model fitting was run using matlab’s fmincon function. For parameters that were not fixed in value the fit lower and upper bounds were set to: *σ*_i_ [0, 20], *σ*_s_ [0, 10], *σ*_a_ [0, 300], bias [-50, 50], lapse [0, 1], lapse bias [0, 1], *α* [-2, 3], *φ* [-3, 0.6], *τ*_*φ*_ [-4, 5], window of integration [1, 1000], *λ*[-5, 5]. Maximum number of iterations was set to 1000, the maximum number of function evaluations was set to 10,000, and the constraint tolerance was set to 10^-7^. The actual values of *φ* and *τ*_*φ*_ were 10^*φ*^ and 10^*τφ*^ respectively. This allowed us to explore the parameter values in logarithmic space. For the burst model the bias parameter assumed the role of burst threshold. Seeds were selected with values in the range [0, 10] and fitting was constrained to remain between [0, 50].

The fitting minimized the least squares difference between the model output probability of making a right response and the subject’s actual response (left = 0 and right = 1). Least squares was used rather than negative log-likelihood as it does not introduce any systematic biases at response probabilities close to 0 and 1. The negative log-likelihood was computed from the best fit parameter values for BIC analysis.

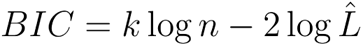

Where k is the number of free parameters in the model, n is the number of trials the model is fit to, and 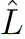 is the maximum likelihood of the data given the best fit parameters.

### Snapshot and Full Integration Model

Below are the equations used to compute the probability of making a right response given a particular stimulus for the snapshot and full integration model. The full integration model is simply the snapshot model with a window of integration duration of 1 s. Before applying click adaptation all clicks are assumed to have a magnitude of 1. Click adaptation is given by the following equation:

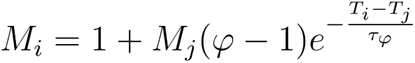

Where M_i_ is the magnitude of the i^th^ click, M_j_ is the magnitude of the j^th^ click, *φ* is the click adaptation scale factor, T_i_ is the time of the i^th^ click, T_j_ is the time of the j^th^ click, and *τ*_*φ*_is the click adaptation time constant, for i > j. When *φ* > 1, clicks facilitate subsequent clicks, and when *φ* < 1 clicks depress subsequent clicks. *φ* and *τ*_*φ*_ are free parameters fit by the model.

After click adaptation is applied and the magnitude of each click is determined, the integrator’s memory time constant is applied to further scale the magnitude of each click:

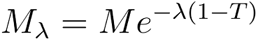

Where M_*λ*_ is the magnitude of the click after applying the integrator’s memory time constant, M is the magnitude of the click before, *λ* is the inverse of the time constant, and T is the time of the click relative to the start of the stimulus. *λ* is a free parameter fit by the model.

To analytically compute the distribution of the accumulated value at the end of the stimulus we separately calculate the mean and standard deviation (std) of the distribution.

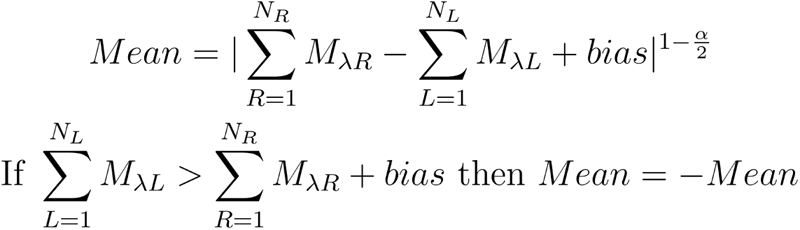

Where M_*λ*R_ (M_*λ*L_) is the magnitude of right (left) favoring clicks computed above, and N_R_ (N_L_) is the number of right (left) favoring clicks that are within the window of integration. Bias and *α* are free parameters fit by the model.

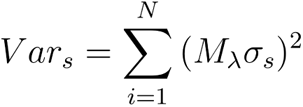

Where N is the total number of clicks. *σ*_s_ is a free parameter fit by the model.

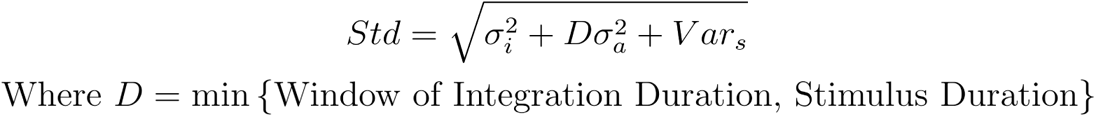

*σ*_i_ and *σ*_a_ are free parameters fit by the model. The window of integration duration is fixed at 1 s in the full integration model and is a free parameter in the snapshot model.

Using the mean and standard deviation of the distribution we calculate the probability of responding right without considering lapse:

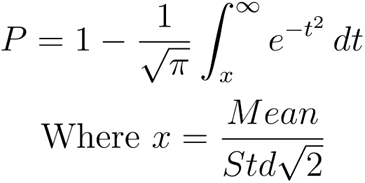

Final we apply the lapse to compute the final probability of responding right:

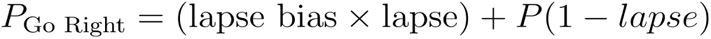

Lapse and lapse bias are free parameters fit by the model.

The probability of making a right response is computed for all placements of the window of integration and an average is taken to compute the probability of responding right for that stimulus.

### Burst Model

Computing the probability of making a right response for the burst model has a few differences from the equations above. *λ* and *σ*_a_ are both fixed at 0 (See Table 1). *P* is only computed for periods where the window of integration samples clicks that all favor the same response. Since the window of integration is sequentially sampling the stimulus, any period that results in a decision commitment prevents subsequent clicks from being sampled. Therefore we cannot simply take a weighted average of all possible positions of the window of integration but rather must scale the probability of committing at a particular position by the probability the model has not already committed to a decision. The probability of committing to a response at the i^th^ click, where P is the probability computed above, is given by:

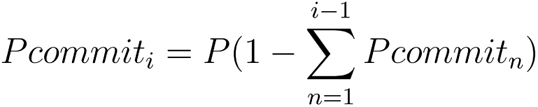

The probability of making a right response (without considering lapse) given a particular stimulus is therefore the sum of Pcommit for all periods when the window of integration samples right clicks plus one half the probability the model does not commit and must guess:

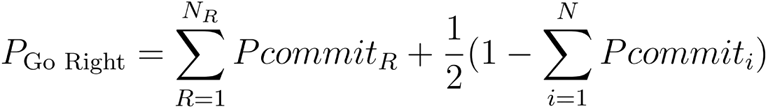

Finally, lapse is applied as shown above for the snapshot and full integration models.

### Single Click Model

Since the single click model does not have click adaptation or a memory time constant for the integrator we skip those calculations and proceed to computing the mean and standard deviation of the accumulator value after sampling a left click, a right click, or no click. All clicks therefore have a magnitude of 1.

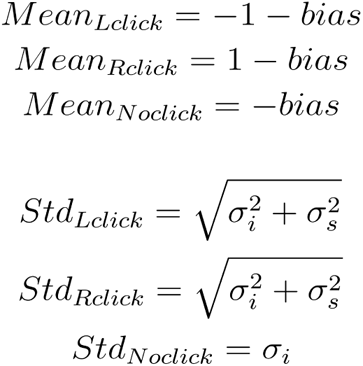

The window of integration start time uniformly samples the entire stimulus period. The model responds to the first click that falls within the window, and the probability of making a right response given that window start time is given by:

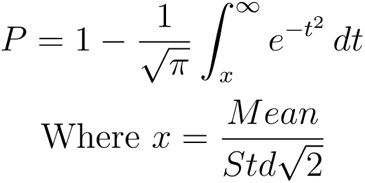

using the appropriate Mean and Std from above. The overall probability of making a right response is the average P across all possible start times for the window of integration. Lapse is applied to P as shown above for the snapshot and full integration model.

### Psychophysical Kernel

To compute the psychophysical kernel the click difference *d* (#R - #L) is first computed within bins (here 100 ms) that tile the stimulus for each trial. All stimuli across trials that are at least 500 ms in duration are aligned to either the stimulus onset or offset. Only clicks that fall within the first 5 bins are considered in this analysis. The total weighted click difference for a trial is given by:

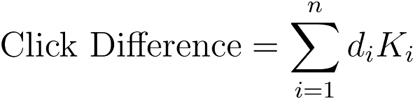

Where i is the bin number, n is the total number of bins (here n = 5), *d* is the click difference within the bin, and *K* is the value of the kernel for that bin (a free parameter that is fit). A psychometric curve is then fit to the rat’s response given the weighted click difference (see equation above, *F* is the fraction “go right”). The probability *P* of the rat’s response given the weighted click difference is simply *F* on trials where the rat responds “right” and 1-*F* on trials where the rat responds “left”. The negative log-likelihood across all trials is given by:

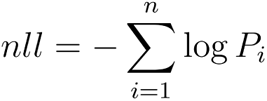

Where i is the trial number, n is the total number of trials, and P is the probability of the rat’s response given the best fit psychometric curve. The psychophysical kernel is the vector *K* that minimizes *nll*.

### The Dataset

The dataset can be downloaded as a single .zip file from zenodo.org (Kopec et al., 2024) DOI 10.5281/zenodo.13352119. Any work that uses this dataset should cite it and this manuscript. Some of the labels in this dataset use the word bup, this means the same as click.

Each file is a matlab .mat file named for the rat that provided the data and contains a single variable, ratdata:

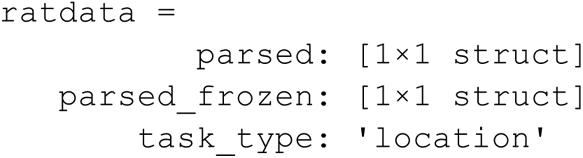

The task type identifies which version of the Poisson Clicks Task the rat trained on, Location or Frequency. The parsed field contains all trail data from unique, i.e. non-frozen trials.

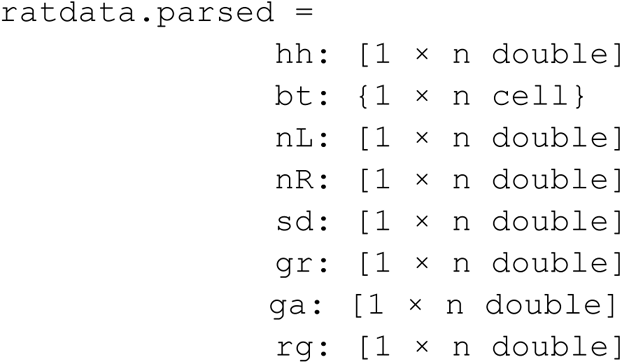

Where n is the number of non-violation non-frozen trials the rat performed.

`hh` - is a vector identifying which trials the rat got correct: 0 if they got it wrong, 1 if they got it correct

`bt` - is a cell which contains information about the stimulus for each trial

`nL` - is a vector identifying how many left evidence clicks each trial contains nR - is a vector identifying how many right evidence clicks each trial contains sd - is a vector of the stimulus duration in seconds for each trial

`gr` - is a vector identifying which response the rat made: 0 if they responded left, 1 if they responded right

`ga` - is a vector of the gamma value used to generate the stimulus on each trial

`rg` - is a vector identifying how the reward was assigned on each trial: 0 if it was offered with the side that played more clicks, 1 if it was offered on the side with the higher generative Poisson click rate

For the rats in this dataset where reward followed the trial’s gamma value, 98.5% of the trials the reward was still offered on the side that played the greater number of clicks.

At a minimum bt is a structure that has two fields:

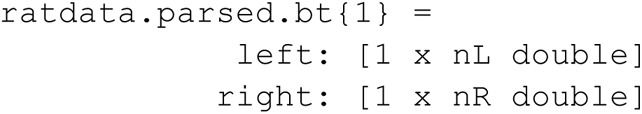

`left` - a vector of the left evidence click times in seconds relative to the stimulus onset

`right` - a vector of the right evidence click times in seconds relative to the stimulus onset

Where nL is the number of left evidence clicks and nR is the number of right evidence clicks. Additional fields can include:

`real_T` - the duration of the stimulus in seconds

`bup_width` - the duration of each click in milliseconds

`base_freq` - the lowest frequency used to make in each click sound in Hertz bup_ramp - defines the taper that smoothes the edges of each click

`first_bup_stereo` - 1 if the first click is a stereo click, 0 if not

`avoid_collisions` - 1 if click trains are selected where clicks do not overlap, 0 if only Poisson statistics are used to select click times

`seed` - the pseudorandom list seed used to generate the stimulus

`is_probe_trial` - 1 if the trial is a “probe” trial. These typically are defined to have a specific stimulus duration, for example 1 second, such that this duration is oversampled. Probe trials exist to facilitate other experiments such as optogenetics, however no trials with optogenetic stimuli are included in this dataset

`is_frozen -` 1 if this is a frozen noise trial, 0 if it is not

`tones` - the specific frequencies used to construct the click in Hertz

In the location version of the task typical settings are:

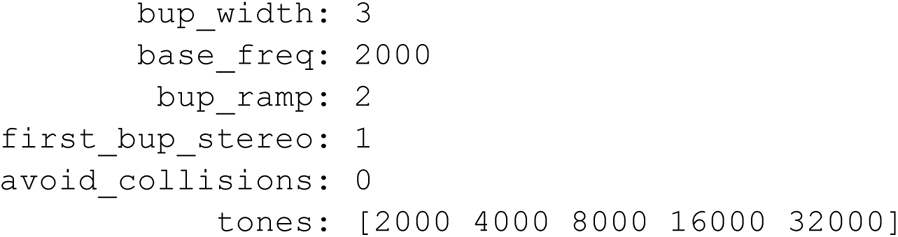

Whereas in the frequency version of the Poisson Clicks Task typical settings that differ are:

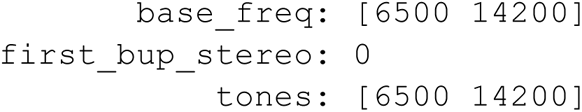

The base frequency and tone fields identify the frequency of the tone used to identify left and right evidence clicks, here 6.5 kHz is the tone frequency for a left evidence click and 14.2 kHz is the tone frequency for a right evidence click. In the frequency task the first click is typically not a stereo click.

The parsed_frozen field contains similar information as the parsed field but for the frozen noise trials:

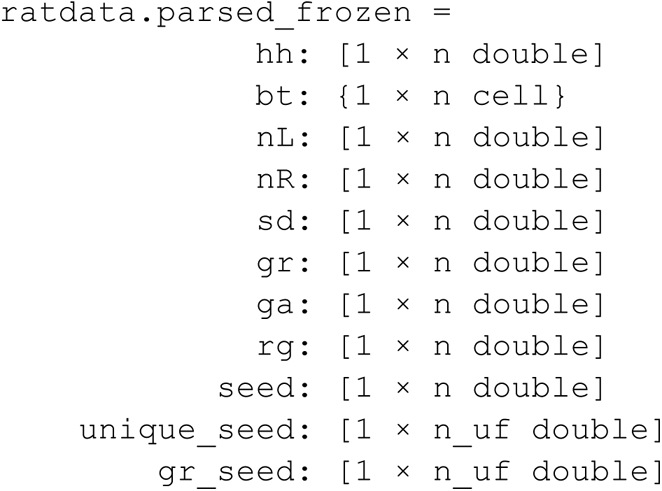

Where n is the number of frozen noise trials the rat performed and n_uf is the number of unique frozen noise trials performed.

`seed` - is a vector of the pseudorandom list seeds used to generate the stimulus for each trial

`unique_seed` - is a vector of the unique seeds used to generate the frozen noise trials

`gr_seed` - is a vector of the fraction of right responses the rat makes given each unique seed

The field names in this dataset are abbreviations:

`hh` - hit history, hit being a trial the subject got correct

`bt` - bup times, bup mean the same as click

`nL` - number of Left nR - number of Right

`sd` - stimulus duration

`gr -` go right, i.e. did the subject respond right

`ga` - gamma

`rg` - reward gamma

## References

Ashwood ZC, Roy NA, Stone IR, International Brain Laboratory, Urai AE, Churchland AK, Pouget A, Pillow JW. 2022. Mice alternate between discrete strategies during perceptual decision-making. Nat Neurosci 25:201–212.

Bogacz R, Brown E, Moehlis J, Holmes P, Cohen JD. 2006. The physics of optimal decision making: a formal analysis of models of performance in two-alternative forced-choice tasks. Psychol Rev 113:700–765.

Bowman NE, Kording KP, Gottfried JA. 2012. Temporal integration of olfactory perceptual evidence in human orbitofrontal cortex. Neuron 75:916–927.

Box GEP. 1976. Science and Statistics. J Am Stat Assoc 71:791–799.

Brody CD, Hanks TD. 2016. Neural underpinnings of the evidence accumulator. Curr Opin Neurobiol 37:149–157.

Bronfman ZZ, Brezis N, Moran R, Tsetsos K, Donner T, Usher M. 2015. Decisions reduce sensitivity to subsequent information. Proc Biol Sci 282. doi:10.1098/rspb.2015.0228

Brunton BW, Botvinick MM, Brody CD. 2013. Rats and humans can optimally accumulate evidence for decision-making. Science 340:95–98.

Busemeyer JR, Townsend JT. 1993. Decision field theory: a dynamic-cognitive approach to decision making in an uncertain environment. Psychol Rev 100:432–459.

Busse L, Ayaz A, Dhruv NT, Katzner S, Saleem A, Schölvinck M, Zaharia AD, Carandini M. 2011. The detection of visual contrast in the behaving mouse. J Neurosci 31:11351–11361.

Calhoun AJ, Pillow JW, Murthy M. 2019. Unsupervised identification of the internal states that shape natural behavior. Nat Neurosci 22:2040–2049.

Carandini M, Churchland AK. 2013. Probing perceptual decisions in rodents. Nat Neurosci 16:824–831.

Cartwright D, Festinger L. 1943. A quantitative theory of decision. Psychol Rev 50:595–621.

Cheadle S, Wyart V, Tsetsos K, Myers N, de Gardelle V, Herce Castañón S, Summerfield C. 2014. Adaptive gain control during human perceptual choice. Neuron 81:1429–1441.

Cisek P, Puskas GA, El-Murr S. 2009. Decisions in changing conditions: the urgency-gating model. J Neurosci 29:11560–11571.

Copernicus N. 1543. De revolutionibus orbium coelestium, libri VI. Johannes Petrejus.

Deverett B, Koay SA, Oostland M, Wang SS-H. 2018. Cerebellar involvement in an evidence-accumulation decision-making task. Elife 7. doi:10.7554/eLife.36781

Ditterich J. 2006. Stochastic models of decisions about motion direction: behavior and physiology. Neural Netw 19:981–1012.

Fechner GT. 1860. Elemente der Psychophysik. Breitkopf u. Härtel.

Gibbon J. 1977. Scalar expectancy theory and Weber’s law in animal timing. Psychol Rev 84:279–325.

Glickman M, Usher M. 2019. Integration to boundary in decisions between numerical sequences. Cognition 193:104022.

Gold JI, Law C-T, Connolly P, Bennur S. 2008. The relative influences of priors and sensory evidence on an oculomotor decision variable during perceptual learning. J Neurophysiol 100:2653–2668.

Gold JI, Shadlen MN. 2007. The neural basis of decision making. Annu Rev Neurosci 30:535–574.

Gold JI, Shadlen MN. 2000. Representation of a perceptual decision in developing oculomotor commands. Nature 404:390–394.

Gupta D, DePasquale B, Kopec CD, Brody CD. 2024. Trial-history biases in evidence accumulation can give rise to apparent lapses in decision-making. Nat Commun 15:662.

Hanks TD, Kopec CD, Brunton BW, Duan CA, Erlich JC, Brody CD. 2015. Distinct relationships of parietal and prefrontal cortices to evidence accumulation. Nature 520:220–223.

Hanks TD, Summerfield C. 2017. Perceptual Decision Making in Rodents, Monkeys, and Humans. Neuron 93:15–31.

Harvey CD, Coen P, Tank DW. 2012. Choice-specific sequences in parietal cortex during a virtual-navigation decision task. Nature 484:62–68.

Hermoso-Mendizabal A, Hyafil A, Rueda-Orozco PE, Jaramillo S, Robbe D, de la Rocha J. 2020. Response outcomes gate the impact of expectations on perceptual decisions. Nat Commun 11:1057.

Huk AC, Shadlen MN. 2005. Neural activity in macaque parietal cortex reflects temporal integration of visual motion signals during perceptual decision making. J Neurosci 25:10420–10436.

Hyafil A, de la Rocha J, Pericas C, Katz LN, Huk AC, Pillow JW. 2023. Temporal integration is a robust feature of perceptual decisions. Elife 12. doi:10.7554/eLife.84045

Katz LN, Yates JL, Pillow JW, Huk AC. 2016. Dissociated functional significance of decision-related activity in the primate dorsal stream. Nature 535:285–288.

Keung W, Hagen TA, Wilson RC. 2020. A divisive model of evidence accumulation explains uneven weighting of evidence over time. Nat Commun 11:2160.

Keung W, Hagen TA, Wilson RC. 2019. Regulation of evidence accumulation by pupil-linked arousal processes. Nat Hum Behav 3:636–645.

Kiani R, Hanks TD, Shadlen MN. 2008. Bounded integration in parietal cortex underlies decisions even when viewing duration is dictated by the environment. J Neurosci 28:3017–3029.

Kiani R, Shadlen MN. 2009. Representation of confidence associated with a decision by neurons in the parietal cortex. Science 324:759–764.

Kopec CD, Luo TZ, Bondy AG, Gupta D, Elliott VA, Charlton JA, Breda JR, Stagnaro WM, Reyes EJ, Sirko AI, Bustos AF, Willock JM, Morrison JM, Osorio KL, Brody CD. 2024. Brody Lab Poisson Clicks Task Dataset Rats 2009 to 2024 Parsed. doi:10.5281/zenodo.13352119

Levi AJ, Yates JL, Huk AC, Katz LN. 2018. Strategic and Dynamic Temporal Weighting for Perceptual Decisions in Humans and Macaques. eNeuro 5. doi:10.1523/ENEURO.0169-18.2018

Levi AJ, Zhao Y, Park IM, Huk AC. 2023. Sensory and Choice Responses in MT Distinct from Motion Encoding. J Neurosci 43:2090–2103.

Meck WH, Church RM. 1983. A mode control model of counting and timing processes. J Exp Psychol Anim Behav Process 9:320–334.

Morcos AS, Harvey CD. 2016. History-dependent variability in population dynamics during evidence accumulation in cortex. Nat Neurosci 19:1672–1681.

Morillon B, Schroeder CE, Wyart V. 2014. Motor contributions to the temporal precision of auditory attention. Nat Commun 5:5255.

Newsome WT, Paré EB. 1988. A selective impairment of motion perception following lesions of the middle temporal visual area (MT). J Neurosci 8:2201–2211.

Nieder A, Miller EK. 2003. Coding of cognitive magnitude: compressed scaling of numerical information in the primate prefrontal cortex. Neuron 37:149–157.

Nienborg H, Cumming BG. 2007. Psychophysically measured task strategy for disparity discrimination is reflected in V2 neurons. Nat Neurosci 10:1608–1614.

Odoemene O, Pisupati S, Nguyen H, Churchland AK. 2018. Visual Evidence Accumulation Guides Decision-Making in Unrestrained Mice. J Neurosci 38:10143–10155.

Palminteri S, Wyart V, Koechlin E. 2017. The Importance of Falsification in Computational Cognitive Modeling. Trends Cogn Sci 21:425–433.

Pardo-Vazquez JL, Castiñeiras-de Saa JR, Valente M, Damião I, Costa T, Vicente MI, Mendonça AG, Mainen ZF, Renart A. 2019. The mechanistic foundation of Weber’s law. Nat Neurosci 22:1493–1502.

Pinto L, Koay SA, Engelhard B, Yoon AM, Deverett B, Thiberge SY, Witten IB, Tank DW, Brody CD. 2018. An Accumulation-of-Evidence Task Using Visual Pulses for Mice Navigating in Virtual Reality. Front Behav Neurosci 12. doi:10.3389/fnbeh.2018.00036

Ptolemy C. 150. The Almagest.

Raposo D, Sheppard J, Schrater P, Churchland A. 2012. Multisensory decision-making in rats and humans. J Neurosci 32:3726–3735.

Ratcliff R. 1978. A theory of memory retrieval. Psychol Rev.

Ratcliff R, McKoon G. 2008. The diffusion decision model: theory and data for two-choice decision tasks. Neural Comput 20:873–922.

Sanders JI, Kepecs A. 2012. Choice ball: a response interface for two-choice psychometric discrimination in head-fixed mice. J Neurophysiol 108:3416–3423.

Schwarz G. 1978. Estimating the Dimension of a Model. Ann Stat 6:461–464.

Scott BB, Constantinople CM, Erlich JC, Tank DW, Brody CD. 2015. Sources of noise during accumulation of evidence in unrestrained and voluntarily head-restrained rats. Elife 4:e11308.

Shadlen MN, Newsome WT. 1996. Motion perception: seeing and deciding. Proc Natl Acad Sci U S A 93:628–633.

Stine GM, Zylberberg A, Ditterich J, Shadlen MN. 2020. Differentiating between integration and non-integration strategies in perceptual decision making. Elife 9:e55365.

Thorndike EL. 1911. Animal intelligence: Experimental studies. New York, The Macmillan company.

Thura D, Beauregard-Racine J, Fradet C-W, Cisek P. 2012. Decision making by urgency gating: theory and experimental support. J Neurophysiol 108:2912–2930.

Uchida N, Kepecs A, Mainen ZF. 2006. Seeing at a glance, smelling in a whiff: rapid forms of perceptual decision making. Nat Rev Neurosci 7:485–491.

Usher M, McClelland JL. 2001. The time course of perceptual choice: the leaky, competing accumulator model. Psychol Rev 108:550–592.

Waskom ML, Kiani R. 2018. Decision Making through Integration of Sensory Evidence at Prolonged Timescales. Curr Biol 28:3850–3856.e9.

Wilson RC, Collins AG. 2019. Ten simple rules for the computational modeling of behavioral data. Elife 8. doi:10.7554/eLife.49547

Wyart V, de Gardelle V, Scholl J, Summerfield C. 2012. Rhythmic fluctuations in evidence accumulation during decision making in the human brain. Neuron 76:847–858.

Yang T, Shadlen MN. 2007. Probabilistic reasoning by neurons. Nature 447:1075–1080.

Yates JL, Park IM, Katz LN, Pillow JW, Huk AC. 2017. Functional dissection of signal and noise in MT and LIP during decision-making. Nat Neurosci 20:1285–1292.

Znamenskiy P, Zador AM. 2013. Corticostriatal neurons in auditory cortex drive decisions during auditory discrimination. Nature 497:482–485.

